# A Unified Comparison of 10X Genomics and Parse Biosciences scRNA-seq Technologies

**DOI:** 10.64898/2026.10.09.758116

**Authors:** Maya Caskey, Ingileif Hallgrimsdottir

## Abstract

Droplet-based and split-pool barcoding are the two dominant high-throughput single-cell RNA-seq paradigms, yet their systematic biases remain incompletely characterized. We compared seven matched 10X Genomics Chromium and Parse Biosciences Evercode WT sample pairs from five studies spanning multiple tissues and species focusing on gene counts and compositional differences rather than cell-type resolution. By separating Parse reads by priming strategy, we distinguished biases arising from priming chemistry from platform-level effects. These effects were largely independent, with platform-level differences substantially larger. Priming strategy introduced a GC-content bias within Parse, whereas the two platforms showed a consistent compositional shift across priming strategies, supported by differential capture of nuclear- and cytoplasmic-enriched transcripts. We interpret this shift as a relative enrichment of nuclear RNA in Parse. Finally, using contamination measures that do not assume a droplet-specific model, we found background RNA contamination in Parse to be comparable in magnitude to that in 10X, contrary to claims of substantially lower contamination. Together, these results indicate that expression measurements and cell-type composition are not directly interchangeable between the two platforms.

## Introduction

Over the past couple of decades, single-cell methods have become essential components of biological discovery, as they are uniquely positioned to capture genome-wide molecular variation at the resolution of individual cells (Tanay and Regev, 2017; Kolodziejczyk et al., 2015). Among the several single-cell RNA-seq experimental paradigms, droplet-based and split-pool barcoding technologies are the highest-throughput techniques, able to profile up to millions of cells at a time (Zheng et al., 2017; Rosenberg et al., 2018). Both technologies use physical or conceptual isolation of cells to label each mRNA molecule with a cell barcode and unique molecular identifier (UMI) through a combination of reverse transcription, ligation, and amplification. The resulting cDNA library can then be sequenced and each read assigned to a single cell and molecule during downstream processing.

Droplet-based technologies, commercialized by 10X Genomics, are the older of the two and currently the most widely used (Zheng et al., 2017). With this set of methods, cells and RNA-barcoded beads are separately loaded onto a microfluidics device, which encapsulates a cell and bead pair within a droplet containing lysis buffer. Once encapsulated, the cell lyses in the buffer, releasing RNA that binds to the bead primers and initiates reverse transcription.

In contrast, split-pool barcoding technologies do not physically separate cells (Rosenberg et al., 2018; Cao et al., 2017). Instead, the cells themselves act as the reaction chamber for RNA priming and reverse transcription, and cells are permeabilized to allow reagents into the cytoplasm. Transcripts are uniquely labelled through multiple barcoding rounds. In each round, pooled cells are divided onto a multi-well plate. In the first round, RNA sequences are captured by a DNA primer containing a well-unique barcode and a UMI, then reverse transcribed. In subsequent rounds, a well-unique barcode is ligated onto the end of a cDNA molecule’s existing primer which is then amplified. In an optional final round, cells are divided into sub-libraries before cell lysis, and the Illumina sample indices added during PCR amplification act as an additional barcode. This approach relies on the stochastic combinatorics of the split-pool process to ensure that each cell is labeled with a unique barcode combination with high probability.

Both technologies are now widely in use, particularly in their commercialized forms, and have been shown to perform similarly with respect to distinguishing cell types. However, they also exhibit systematic biases that complicate direct comparison and generalization across platforms. Several points of technical difference between the Parse Biosciences and 10X Genomics kits could plausibly contribute to such biases. The Chromium 3’ kit profiles the contents of live, lysed cells, whereas the Evercode WT kit profiles the contents of formaldehyde-fixed and permeabilized cells. The two encapsulation paradigms also differ in their susceptibility to doublets, barcode collisions, and ambient RNA contamination, all of which are known to distort single-cell data. Finally, the platforms differ in RNA priming strategy: the 10X Chromium kit relies exclusively on oligo(dT) priming of the poly(A) tail common to most mRNA, whereas the Parse Evercode WT kit combines oligo(dT) priming with random hexamer priming, which additionally captures non-polyadenylated RNA species such as certain long non-coding RNAs.

Several studies from different groups, spanning multiple species and tissues, have characterized aspects of these biases, particularly between the 10X Genomics Single-Cell 3’ Chromium kit and the Parse Biosciences Evercode WT kit (Gezelius et al., 2024; Xie et al., 2024; De Simone et al., 2025; Filippov et al., 2024; Hornung et al., 2023; King et al., 2026; Brown et al., 2024; Sarieva et al., 2026). These studies are difficult to reconcile with one another, as each examines a different subset of features, and none draws on a sufficient number of biological and technical replicates across tissues and species to support a general conclusion. No study to date has attempted to unify these findings into a single, direct comparison.

In this paper, we compare seven matched 10X Genomics Chromium and Parse Biosciences Evercode WT sample pairs spanning multiple tissues, species, and kit versions. Within Parse, we separate oligo(dT)-primed reads from random-hexamer-primed reads to determine the extent to which systematic biases originate from priming strategy rather than from other platform differences. In addition to library-level quality metrics, we perform bulk differential expression and gene ontology analysis to characterize compositional differences in gene capture both between platforms and between priming strategies within Parse. We find that 10X and Parse diverge primarily along an axis consistent with differential nuclear RNA capture, a bias that is largely independent of priming strategy and reproducible across species; we additionally find a smaller, distinct bias between priming strategies within Parse that is best explained by transcript GC content.

## Results

To characterize systematic differences between droplet-based and split-pool single-cell RNA-seq platforms, we compared seven matched 10X Genomics Chromium and Parse Biosciences Evercode WT sample pairs drawn from five previously published studies (Table 1). The datasets span three tissue types (cell lines, peripheral blood mononuclear cells (PBMCs), and thymocytes) and three species contexts (human, mouse, and human-mouse mixture). For each experiment, Parse reads were further partitioned by priming strategy into three constituent datasets: reads captured by oligo(dT) priming alone (“Parse 3”‘), reads captured by random hexamer priming alone (“Parse randO”), and all Parse reads together (“Parse combined”). Each of the four datasets in a given experiment – 10X, Parse 3’, Parse randO, and Parse combined – was independently subsampled from its own respective pool of reads to the depth of the smallest dataset in that experiment (Figure S1.). This allows for direct comparison of library quality and composition at matched sequencing depth without conflating depth differences with platform or priming differences.

**Table 1.** Publicly available datasets analyzed. In this paper, all datasets are referred to by their ID. Note that in Exp 4a and Exp 4b, 10X reads are subsampled from the same library, and for Exp 2a and Exp 2b, Parse samples were multiplexed on the same plate.

| ID | Species | Tissue | Subsampled Read Depth | 10X Chromium Single Cell 3' Kit | Parse Evercode WT Kit | Publication |
| --- | --- | --- | --- | --- | --- | --- |
| Exp 1 | *H. sapiens* | REH Cell Line | 164,834,585 | v3 (2018) + MULTI-seq | Mini v1 | Gezelius et al. (2024) |
| Exp 2a | *H. sapiens* | PBMCs | 89,116,454 | v3 (2018) | v2 (2022) | Xie et al. (2024) |
| Exp 2b | *H. sapiens* | PBMCs | 98,133,317 | v3 (2018) | v2 (2022) | Xie et al. (2024) |
| Exp 3 | *H. sapiens* | PBMCs | 64,554,749 | v3.1 (2019) | v2 (2022) | De Simone et al. (2025) |
| Exp 4a | *M. musculus* | Thymocytes | 639,108,133 | v3 (2018) + TotalSeq-B | v2 (2022) | Filippov et al. (2024) |
| Exp 4b | *M. musculus* | Thymocytes | 188,781,476 | v3 (2018) + TotalSeq-B | Mini v2 (2022) | Filippov et al. (2024) |
| Exp 5 | *H. sapiens* / *M. musculus* | K562 / mESC<br>Barnyard | 14,026,910 | v3 (2018) | v2 (2022) | Hornung et al. (2023) |

### 10X sequencing results in higher library quality regardless of priming strategy though differences in gene diversity are less clear

10X libraries were of higher quality than Parse libraries by several measures, regardless of priming strategy. When aligned to either the transcriptome with kallisto (Bray et al., 2016) (Fig. 1A and 1B) or the genome with STAR (Dobin et al., 2013) (Fig. S2A), the 10X library had a lower percentage of unmapped reads and a higher percentage of valid barcodes than the matching Parse libraries. In addition, when reads were aligned to the genome with STAR, 10X libraries yielded a greater number of uniquely mapped reads than any of the three Parse datasets (Fig. S2B). Within Parse, the randO dataset consistently produced the fewest uniquely mapped reads of the three subsampled datasets.

**Fig 1.**
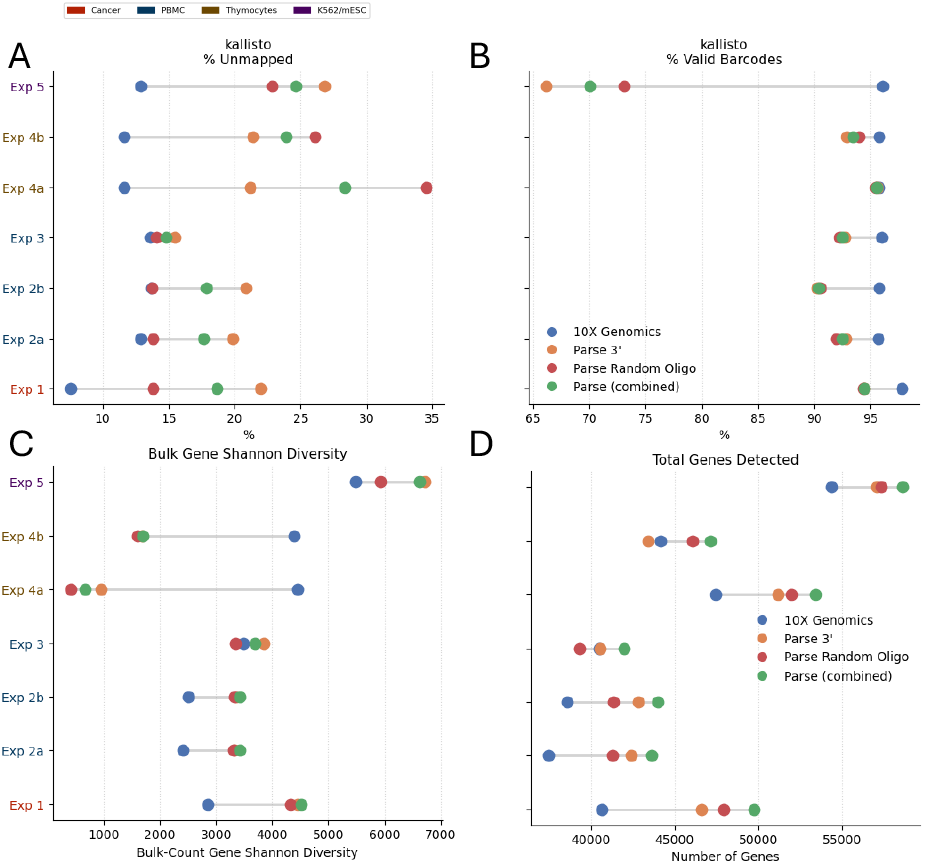
10X exhibits consistently higher library quality than Parse, with less consistent differences in gene diversity. Dot plots showing the range of each kallisto pseudoalignment metric across the subsampled datasets from each experiment: (A) percent of reads unmapped by kallisto; (B) percent of valid barcode combinations recovered by kallisto. Two diversity metrics computed on unfiltered gene counts: (C) shannon entropy over total counts per gene; (D) total genes detected. Experiment labels in (A) and (C) are colored by tissue.

Several of the studies from which these datasets were drawn report that Parse captures a greater diversity of transcripts than 10X (Filippov et al., 2024; Xie et al., 2024; Brown et al., 2024; Gezelius et al., 2024). Sarieva et al. (2026) is the only study to report a greater number of genes captured by 10X, though we did not analyze their data ourselves. Our results generally support this claim for the majority of experiments analyzed. For almost all experiments, both Shannon entropy (Fig. 1C) and total genes detected (Fig. 1D) were lower for 10X than for Parse combined, with the exception of Experiment 4a and 4b. In Experiment 4a and 4b, 10X showed substantially higher Shannon entropy than all Parse datasets despite detecting fewer total genes, except in Experiment 4b, where 10X detected more genes than Parse 3’. In addition, in Experiment 3, 10X detected more genes than Parse randO.

Across all experiments, the Parse combined dataset detected more genes than either the Parse 3’ or Parse randO datasets alone, consistent with the two priming strategies capturing partially non-overlapping sets of transcripts. We note, however, that because Parse combined is independently sub-sampled from the full pool of Parse reads rather than constructed as the union of the 3’ and randO datasets, this result is also consistent in part with the combined pool simply being more diverse prior to subsampling.

### Variance between 10X and Parse datasets is dominated by batch effect between platforms

Across all experiments, 10X and Parse datasets showed high concordance (Fig. 2A), as expected for matched samples profiling the same underlying biology: the concordance correlation coefficient (CCC) exceeded 0.8 between 10X and each of the three Parse datasets. However, the variance attributable to platform substantially exceeded the variance attributable to priming strategy within Parse. Across all comparisons, CCC values between any two of the three Parse datasets (3’, randO, and combined) were consistently higher than CCC values between 10X and any individual Parse dataset.

This pattern was also evident in a principal component analysis of the filtered, shifted-CLR normalized (Booeshaghi et al., 2026) bulk gene counts across all human datasets (Fig. 2B). PC1 (48.9% of variance) separated PBMC samples from cancer cell line samples, reflecting expected biological differences between sample types. PC2 (23.8% of variance) separated samples by platform, with 10X datasets taking positive values and all three Parse datasets taking negative values. Within a given experiment, Parse 3’ clustered more closely with the Parse randO and Parse combined datasets on PC2 than with 10X, indicating that the platform-driven batch effect dominates over both priming strategy and experiment-of-origin effects within Parse. Parse 3’ datasets were, however, consistently shifted toward more positive PC2 values relative to Parse randO and Parse combined.

**Fig 2.**
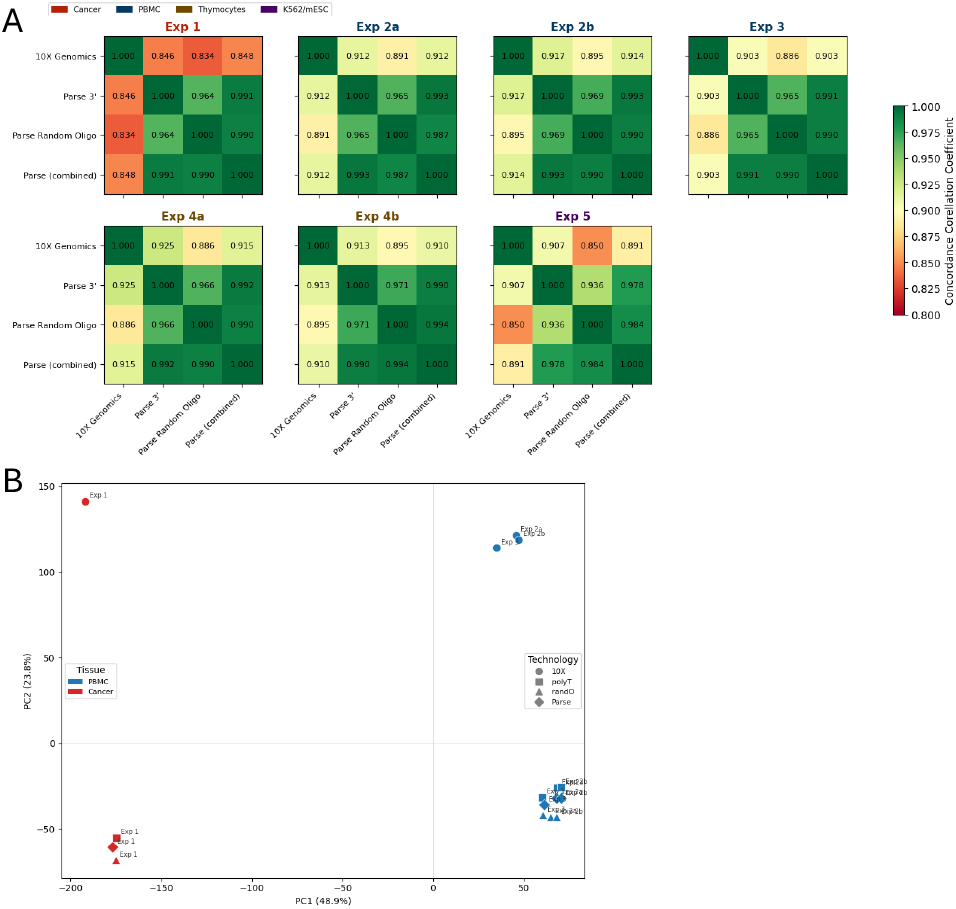
Platform contributes more variance between datasets than priming strategy. (A) PCA of unfiltered and shifted-CLR normalized bulk counts for each subsampled dataset from the four human experiments. (B) Heatmaps of the concordance correlation coefficient (CCC) between filtered and logCPM normalized bulk counts for the four datasets within each experiment. Experiment labels are colored by tissue as in Fig. 1.

Consistent with 10X and Parse 3’ sharing a common priming strategy, CCC values between 10X and Parse 3’ were consistently higher than CCC values between 10X and Parse randO across experiments. These results suggest that the primary source of variation between 10X and Parse is due to platform-level batch effect rather than priming strategies, while differences in priming strategy contribute to batch effect both within and between platforms but to a lesser extent.

### Compositional differences between 10X and Parse are consistent with differential nuclear RNA capture

Beyond the platform-level batch effect described above, 10X and Parse datasets differed in the relative abundance of several transcript classes (Fig. 3), with Parse exhibiting a bias towards nuclear transcripts. The nucleus is known to contain a greater number of unprocessed transcripts, and within each experiment, the mean percentage of nascent (unspliced) counts per cell was higher in Parse than in 10X. Across experiments, the mean nascent percentage in Parse ranged from 35.88% to 59.85%, compared to 13.81% to 40.70% in 10X, with the Parse 3’ and randO datasets closely tracking the combined Parse range.

**Fig 3.**
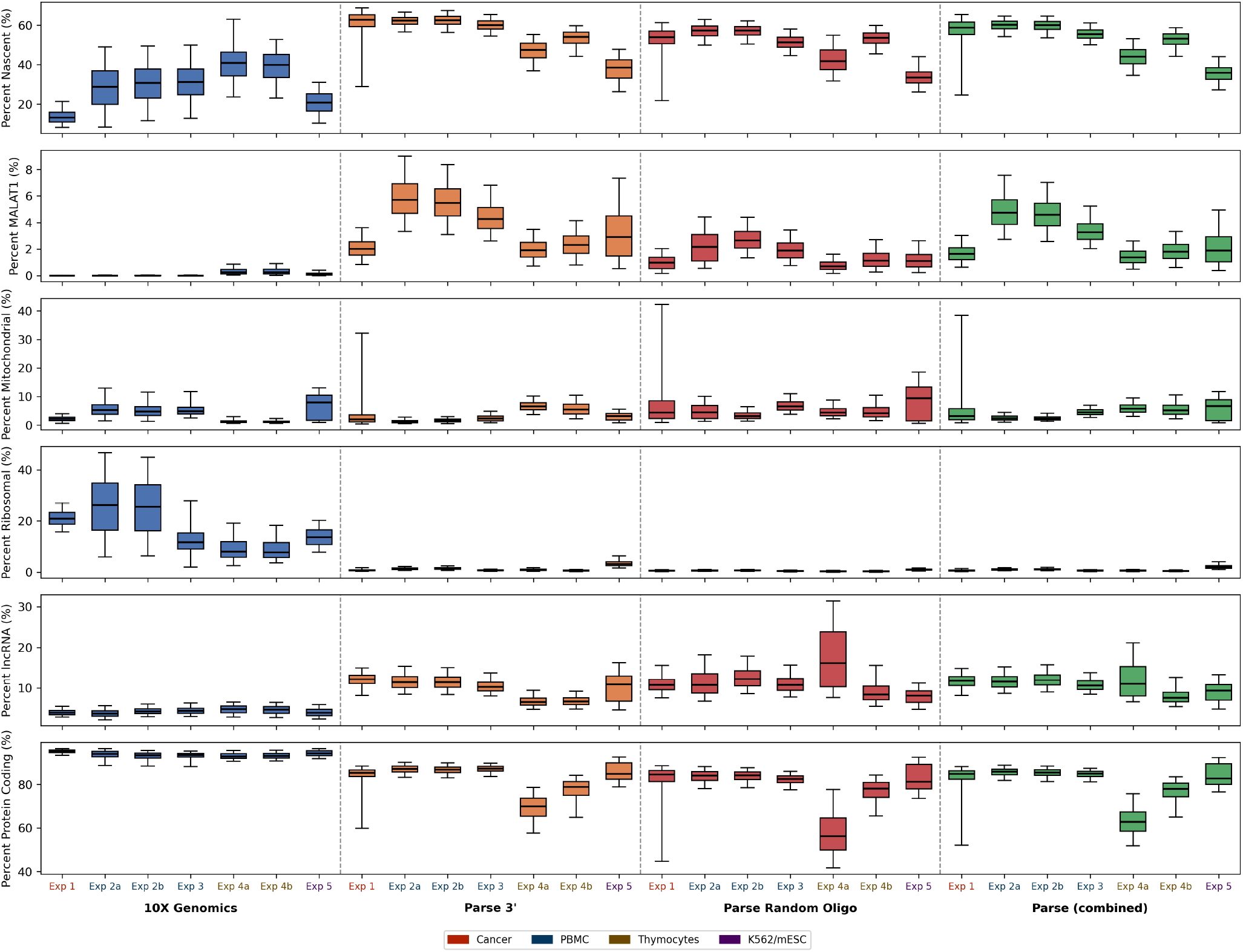
Compositional differences between 10X and Parse are consistent with differential nuclear RNA capture. For all experiments and sub-sampled datasets, box plots of the percentage of filtered counts per cell attributable to each of six categories: nascent (unspliced) counts as assigned by kallisto; MALAT1; mitochondrial genes (prefix MT-); ribosomal protein genes (RPS/RPL and MRPS/MRPL); lncRNA; and protein-coding genes. Outliers beyond the 5th and 95th percentiles are not shown here. Experiment labels are colored by tissue as in Fig. 1.

This pattern extended to individual transcripts and to broader RNA classes, and in two cases the platforms separate almost entirely. MALAT1, a transcript retained almost exclusively in the nucleus (Clarke and Bader, 2024), was substantially depleted in 10X relative to Parse: the mean per-cell MALAT1 percentage across 10X datasets ranged from <0.01% to 0.33%, compared to a mean of 1.44% to 4.87% in Parse, with Parse 3’ and randO again tracking the combined Parse range. Conversely, ribosomal protein-coding RNA was depleted in Parse relative to 10X: mean per-cell ribosomal percentage in Parse ranged from 0.44% to 2.15%, compared to a mean of 9.00% to 25.96% in 10X, a trend which was also noted in King et al. (2026) and Sarieva et al. (2026). A similar pattern of depletion held at the level of broader RNA classes: mean per-cell protein-coding percentage ranged from 92.79% to 95.26% in 10X compared to 63.05% to 85.60% in Parse, while mean per-cell long non-coding RNA (lncRNA) percentage ranged from 8.09% to 12.16% in Parse compared to only 3.81% to 4.75% in 10X, a relationship which was also found in Sarieva et al. (2026).

These results are consistent with a shift toward nuclear RNA capture in Parse relative to 10X. The intronic fraction and MALAT1 percentage are both known to be associated with higher nuclear capture (Clarke and Bader, 2024; Muskovic and Powell, 2021). The shift we observe in Parse toward higher MALAT1 percentage, higher intronic read percentage, lower protein-coding percentage, and higher lncRNA percentage parallels differences previously reported between single-nucleus and single-cell RNA-seq, where single-nucleus protocols show the same directional shifts relative to single-cell protocols (Selewa et al., 2020).

In contrast to this nuclear-capture bias, 10X and Parse showed no consistent difference in indicators of cellular damage. Mean per-cell mitochondrial count percentage ranged from 2% to 8% across both platforms and all experiments, with no clear trend distinguishing 10X from Parse; an increase in mitochondrial RNA percentage is generally taken as an indicator of lower cell quality, as the mitochondrial membrane is more resistant to disruption than the plasma membrane and so mitochondrial contents are less likely to be released prior to lysis (Ilicic et al., 2016). The absence of a platform-level difference in mitochondrial percentage suggests that the nuclear-capture bias observed between 10X and Parse reflects a difference in nuclear RNA retention specifically, rather than a general difference in cell quality or damage between the two protocols.

### Differential expression analysis reveals broad shifts in compositional gene expression between technologies

To further characterize the platform-level bias described above, we performed differential expression analysis with edgePython on the bulk counts from the three human PBMC datasets, blocking on study of origin. Because 10X and Parse 3’ share a common priming strategy, we restricted this comparison to 10X versus Parse 3’ to isolate platform-level effects from priming-level effects.

Despite sharing a priming strategy, 10X and Parse 3’ were readily separable in an MDS plot of the bulk counts (Fig. S3A). Dimension 1 (74.8% of variance) separated 10X from Parse 3’, indicating that platform is the dominant axis of variation even when priming strategy is held constant.

Bulk differential expression analysis identified substantial compositional shifts between platforms: at an FDR threshold of 0.01, 6,346 genes were significantly enriched in 10X and 5,418 in Parse 3’ (Fig. 4A). As a cros-sspecies validation, we asked whether the genes found to be platform-enriched in the human PBMC comparison showed consistent enrichment in an independent pair of mouse thymocyte experiments (Experiments 4a (Fig. 4D) and 4b (Fig. S4A)), which were not part of the original PBMC analysis. For genes significantly enriched in 10X in the PBMC comparison (FDR *<* 0.01), the mouse orthologs showed higher bulk logCPM in 10X than in Parse 3’: 88.46% in Experiment 4a, and 86.57 in Experiment 4b. The converse held only weakly: for genes enriched in Parse in the PBMC comparison, the orthologs favored Parse in just 61.56% of cases in Experiment 4a and 70.97% in Experiment 4b.

**Fig 4.**
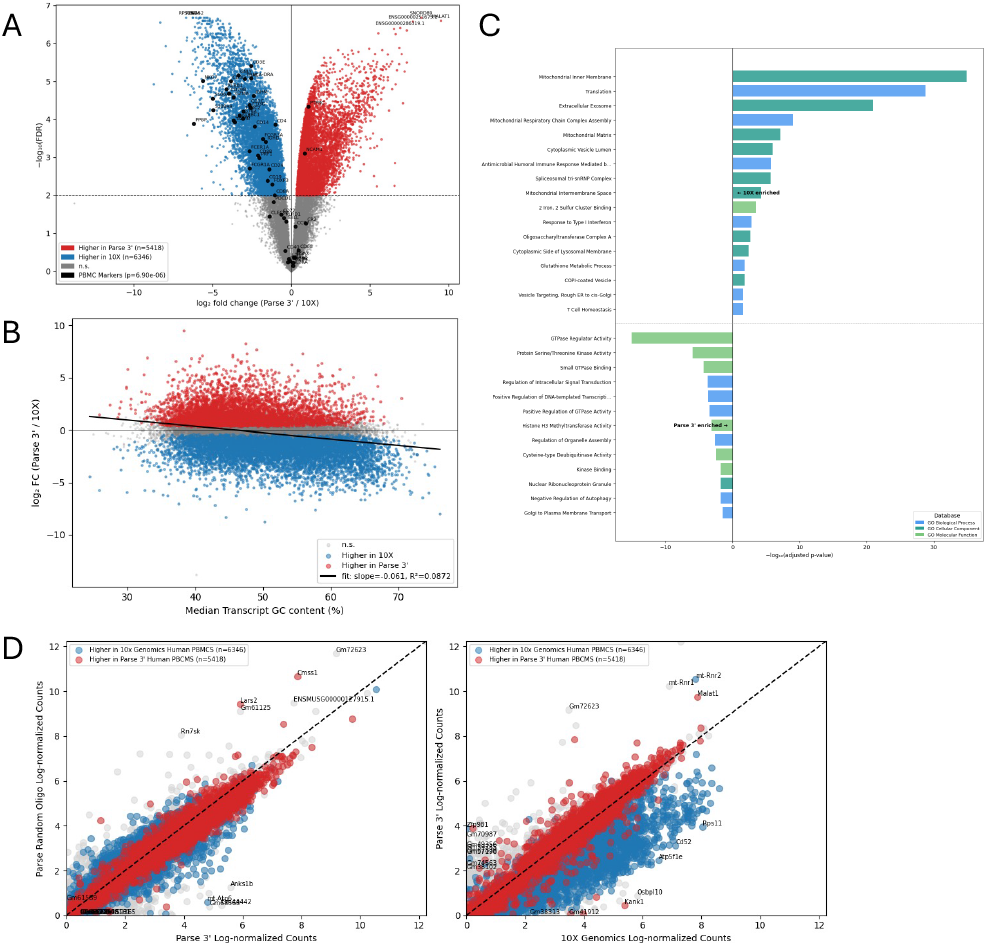
Differential expression analysis reveals broad shifts in compositional gene expression between technologies. Bulk edgePython analysis comparing 10X to Parse 3’ across the three human PBMC experiments, blocking on study of origin. Volcano plot at an FDR threshold of 0.01; significant genes enriched in 10X labelled in blue and enriched in Parse labelled in red, canonical PBMC markers labelled in black, and the five most significant genes labelled. (B) Median Transcript GC content against log-fold change (Parse 3’/10X), with line of best fit. Significant genes enriched in 10X are labeled in blue and enriched in Parse labelled in red. (C) GO term analysis (goseq) controlling for GC content, showing ^*−* log_10_^ adjusted p-values for the most representative terms enriched in 10X and in Parse 3’ that pass the 0.05 adjusted p-value threshold. (D) On the Exp 4a mouse thymocyte datasets: filtered logCPM bulk gene counts compared between Parse randO and Parse 3’ and Parse 3’ and 10X. Genes are colored by whether the ortholog was significantly differentially expressed (FDR < 0.01) in favor of 10X (blue) or Parse 3’ (red); genes lacking orthologs or not significantly differentially expressed are shown in grey.

Because RNA-seq methods are known to exhibit GC-content and length biases through several mechanisms, we asked whether either contributed to the differential expression observed between 10X and Parse 3’. Both GC content (Fig. 4B) and median transcript length (Fig. S3B) were significantly associated with platform-level logFC, but the effect sizes were small (*R*^2^ = 0.0872 and 0.0101 respectively), indicating that neither is a primary driver of the platform difference.

Gene ontology enrichment analysis (adj. p *<* 0.05, goseq (Young et al., 2010)) showed that genes enriched in 10X fell into coherent, well-defined categories, whereas genes enriched in Parse 3’ mapped onto a smaller and less coherent set of terms. Because GC content contributed weakly to differential expression, we corrected for it using the Wallenius noncentral hypergeometric test. Of the terms significantly enriched in 10X, 4 were GC-driven and 78 GC-robust; of the terms enriched in Parse 3’, 23 were GC-robust and 26 were GC-driven (Fig. S3C). As many terms were redundant or related, we collapsed those with overlapping gene sets into 22 representative terms enriched in 10X and 2 in Parse 3’ (Fig. 4C).

Genes enriched in 10X were most strongly associated with the mitochondrial inner membrane (GO:0005743, adj. p ~ 10^*−*44^), reflecting the oxidative phosphorylation complexes (NDUF, COX, UQCR, ATP5/6, and SDH families) together with associated assembly factors, import machinery, and mitochondrial ribosomal proteins. This is in line with a previous finding that genes responsible for mitochondrial respiration are enriched in 10X (Sarieva et al., 2026). The next most significant representative term, Translation (GO:0006412, adj. p~ 10^*−*29^) was driven primarily by ribosomal protein-coding genes and translation elongation factors.

Genes encoding the 26S proteasome were likewise depleted in Parse, though captured only indirectly, through extracellular exosome (GO:0070062, adj. p ~ 10^*−*21^), mitochondrial matrix (GO:0005759, adj. p ~ 10^*−*8^), cytoplasmic vesicle lumen (GO:0060205, adj. p ~ 10^*−*6^). Spliceosomal components were also captured by spliceosomal tri-snRP complex (GO:0097526, adj. p ~ 10^*−*6^).

To confirm this pattern of depletion across these abundant protein complexes, we quantified the per-cell percent-count distribution of each gene set (Fig. 5). Across all Parse datasets, the nuclearencoded core OXPHOS genes— spanning complexes I through V (NDUF, SDH, UQCR, COX, and ATP5 families)—were depleted in Parse relative to 10X: mean OXPHOS percentage per cell ranged from 1.10% to 3.76% in 10X against 0.17% to 0.67% in Parse. The 26S proteasome genes (PSMA/B/C/D families) and the spliceosomal SNRP genes (SNRPB/D/E/F/G) followed the same trend, with a lower mean per-cell percentage in every Parse dataset than in its matching 10X dataset across all experiments.

**Fig 5.**
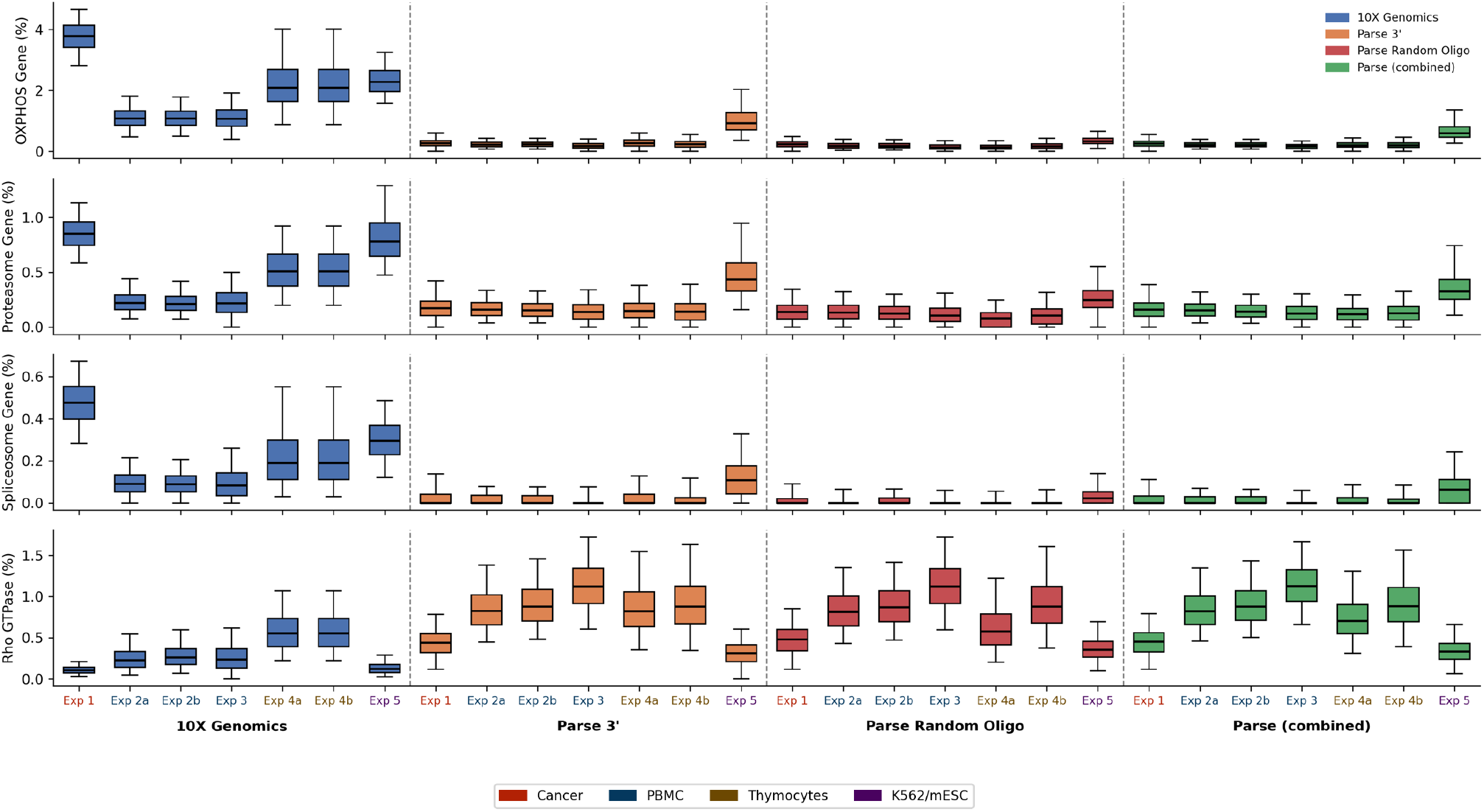
Sets of genes encoding large, ubiquitous protein complexes are depleted in Parse. For all experiments and sub-sampled datasets, box plots of the percentage of filtered counts per cell attributable to each of three categories: oxidative phosporylation pathway (OXPHOS) genes (with prefixes NDUF, UQCR, SDH, COX, or ATP5); proteasome complex genes (with prefixes PSMA/B/C/D); spliceosome complex genes (with prefixes SNRPB/D/E/F/G); Rho GTPase genes (with prefixes DOCK, ARHGAP, and ARHGEF). Outliers beyond the 5th and 95th percentiles are not shown here. Experiment labels are colored by tissue as in Fig. 1.

Although 10X-enriched genes were concentrated in abundant, broadly expressed categories, this extended only weakly to constitutively transcribed housekeeping genes. Using a curated human housekeeping gene set (Hounkpe et al., 2021), 63.1% of housekeeping genes had a negative logFC (favoring 10X), against 51% of genes overall (Fig. S5A).

In contrast to 10X, Parse 3’ was enriched, generally with weaker significance, for genes encoding regulatory proteins rather than large, ubiquitous protein complexes (Fig. 4C). The GO enrichment terms clustered around three related themes: regulation of small-GTPase signaling, protein kinase activity, and chromatin and transcriptional regulation. To confirm this pattern independently of the GO annotation and across all experiments, we quantified the per-cell percent-count distribution of Rho GTPase regulator genes (prefixes DOCK, ARHGAP, and ARHGEF) in each dataset. Consistent with the differential expression result, the mean per-cell percentage of these genes was higher in every Parse dataset than in the matching 10X dataset, across all experiments.

In addition to the findings above, canonical PBMC marker genes also showed greater representation in 10X than in Parse (Fig. 4A). Most canonical markers (82%) had a negative Parse 3’/10X log-fold change, and among the subset that were highly differentially expressed (68%), all but two were enriched in 10X.

In order to confirm whether differentially expressed genes arose from differential transcript capture, as we suspect, rather than differential cell-type composition, we quantified the relative proportion of each cell type within each dataset following cell-type identification with CellTypist (Xu et al., 2023; Conde et al., 2022). Cell-type proportions were broadly similar between platforms within each comparison (Fig. S6). For Experiments 2a and 2b, proportions for 10X and Parse agreed within 3 percentage points across cell types. For Experiment 3, the difference was somewhat larger, with 10X showing evidence of megakaryocyte/platelet contamination (2.1% of cells, versus 0% in Parse); however, megakaryocytes are a small population and not the cell type associated with the markers discussed below, and T-cell proportions in this experiment differed by only 3.36% between platforms. These compositional differences, at most a few percentage points, are too small to account for log_2_ fold-changes of 3–6 (i.e., roughly 10-to 50-fold on a linear scale) observed between platforms for canonical immune markers in the edgePython comparison, such as the cytotoxic marker

NKG7 (log_2_FC(Parse 3’/10X) = *−* 5.62) or the pan-T-cell markers CD3E and CD3D (log_2_FC = *−* 2.55 and *−* 3.84, respectively).

### Differential expression analysis reveals smaller but significant differences between oligo(dT) and randO priming

We next performed the same edgePython differential expression analysis, blocking on study of origin, comparing Parse 3’ to Parse randO reads directly, in order to isolate the effect of priming strategy independent of platform.

Unlike the 10X and Parse 3’ comparison, the Parse 3’ and randO datasets were not separated along the first dimension, accounting for 78.5% of the variance of the MDS plot (Fig. 6A), though they separated along the second dimension accounting for (18.2 %). This suggests that batch effects between experiments were larger than priming bias. Between priming strategies, 5,828 genes were significantly enriched in Parse randO and 3,984 were significantly enriched in Parse 3’ (Fig. 6B). Unlike the 10X versus Parse 3’ comparison, Parse 3’ and Parse randO did not appear to be specifically enriched for PBMC marker genes: only 54% of markers showed a positive log-fold change (randO/3’).

As with the 10X vs. Parse 3’ analysis, we performed cross-validation of differentially expressed genes between Parse 3’ and Parse randO using the mouse thymocyte datasets (Exp 4a and 4b), comparing bulk gene-level logCPM percentages (Fig. 6E and Fig. S4B respectively). This analysis revealed strong concordance with the differential expression results. Of the differentially expressed orthologs enriched in Parse 3’, 93% and 84% had a higher logCPM percentage in Parse 3’ than Parse randO in Experiments 4a and 4b, respectively. Of the differentially expressed orthologs enriched in Parse randO, 73% and 86% had a higher logCPM percentage in Parse randO than Parse 3’ in Experiments 4a and 4b, respectively.

Random hexamer primers bind along the full length of a transcript rather than only at the poly(A) tail, raising the possibility that randO priming preferentially captures longer transcripts. We found instead that the structural bias between Parse 3’ and randO reads was driven primarily by GC content rather than transcript length. In a regression of priming-strategy enrichment on these two variables, gene length explained very little variance (*R*^2^ = 0.0118) compared to GC content (*R*^2^ = 0.2997) (Fig. S7A and Fig. 6C respectively). This bias was also evident when comparing the logCPM gene percentages in the mouse thymocyte datasets (Fig. S4C and S4D).

**Fig 6.**
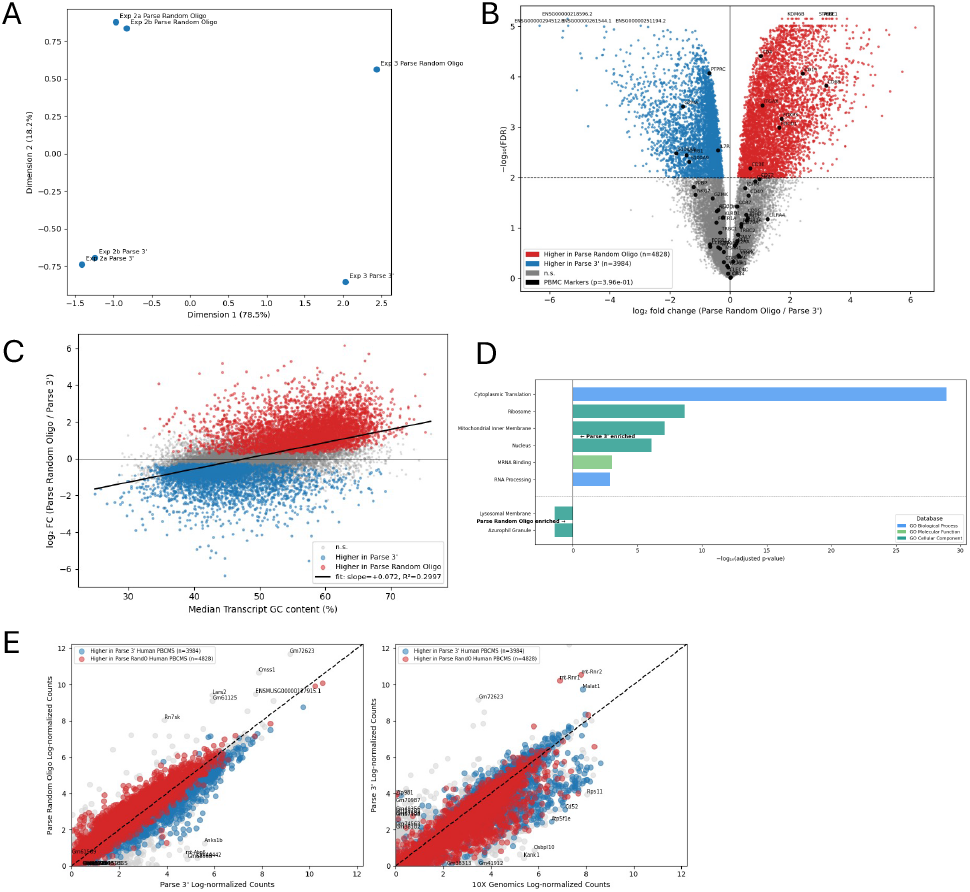
Differential expression analysis reveals smaller but significant differences between oligo(dT) and randO priming. Bulk edgePython analysis comparing Parse 3’ to Parse randO across the three human PBMC experiments, blocking on study of origin. (A) MDS plot. (B) Volcano plot at an FDR threshold of 0.01; significant genes enriched in Parse 3’ labelled in blue and enriched in Parse randO labelled in red, canonical PBMC markers labelled in black, and the five most significant genes labelled. (C) GC content against log-fold change (randO/3’), with line of best fit. (D) On the Exp 4a mouse thymocyte datasets: filtered logCPM bulk gene percentages compared between Parse randO and Parse 3’ and Parse 3’ and 10X. Genes are colored by whether the ortholog was significantly differentially expressed (FDR < 0.01) in favor of Parse 3’ (blue) or Parse randO (red); genes lacking orthologs or not significantly differentially expressed are shown in grey.

We considered whether this GC-content bias could instead be explained by internal off-target priming of oligo(dT) primers at internal adenine-rich regions, rather than by a genuine preference of random hexamers for GC-rich sequence. Gene body coverage plots showed that Parse 3’ reads, while preferentially capturing the 3’ end of transcripts as expected, also captured internal sequence to some degree (Fig. S8). Annotating genes by the number and length of internal adenine single-nucleotide repeats (Svoboda et al., 2022) yielded a weaker *R*^2^ of 0.1889 in the same regression framework, and this association appeared to be driven primarily by the subset of genes differentially enriched in Parse 3’ (Fig. S7B).

Contrary to the expectation that random hexamer priming would be enriched for transcripts lacking a poly(A) tail, this was not consistently the case; enrichment instead tracked more directly with GC content. As an illustrative example, MALAT1 – a transcript lacking a canonical poly(A) tail – was significantly enriched in Parse 3’ rather than randO (logFC(randO/3’) = −1.24, FDR = 2.4e-5), consistent with its relatively low median transcript GC content of 38% despite the absence of a poly(A) tail. NEAT1, another non-polyadenylated transcript, was significantly enriched in randO (logFC = 0.757, FDR = 5.4e-4) and had a higher median transcript GC content of 49%, further supporting the GC-content association over presence or absence of a poly(A) tail as the driver of priming-strategy enrichment.

Enrichment analysis controlling for GC content revealed additional bias not fully explained by GC content alone. Using goseq, we found that of 82 naively significant terms enriched in Parse 3’, 54 were GC-robust, whereas of the 18 naively significant terms enriched in Parse randO, only 2 were GC-robust (Fig. S7C/D). Collapsing redundant terms left 6 representative terms enriched in Parse 3’(Fig.6D). The three most significant Parse 3’-enriched terms— Cytoplasmic Translation (GO:0002181, adj. p ~ 10^*−*29^), Ribosome (adj. p ~ 10^*−*9^), and Mitochondrial Inner Membrane (GO:0005743, adj. p ~ 10^*−*8^)—were driven largely by ribosomal and mitochondrial ribosomal protein genes, transcriptional initiation factors, and mitochondrial respiration genes, similar to the functional gene classes enriched in 10X relative to Parse 3’ in the platform-level comparison above. The randO-enriched terms—lysosomal membrane (GO:0005765, adj. p = 0.039) and azurophil granule (GO:0042583, adj. p = 0.040) —-were only weakly significant in comparison.

As above, we performed cell-typing of the Parse 3’ and Parse randO to confirm that these shifts do not arise from difference in cell composition. Indeed, across all human PBMC experiments, cell-type percentages were consistently within one percentage point of each other across datasets. This suggests that while priming strategy does influence transcript capture, it does not result in compositional bias.

### Parse exhibits substantial markers of ambient RNA contamination, inconsistent with claims of reduced contamination relative to 10X

Ambient RNA contamination is well characterized in droplet-based data, where several methods exist for its estimation and removal. Its extent in split-pool data is less settled, and published estimates disagree in both magnitude and mechanism (Parse Biosciences, 2026; King et al., 2026; Filippov et al., 2024). Because the available estimators have not been validated on split-pool data, we chose not to include them in this analysis. Instead, we rely on quality control metrics and ground-truth data from human-mouse mixtures.

The most direct of these is the composition of barcode space itself. Parse libraries recovered nearly all valid barcode combinations, far exceeding the number of cells loaded. Four of seven experiments recovered greater than 99.9% of possible combinations before filtering; the remaining three (Experiments 3, 4a, and 4b) recovered 97.5%, 90.8%, and 93.9% respectively. Barcode counts fell sharply after filtering, approaching the number of cells loaded, indicating that most recovered combinations correspond to low-count background rather than cells.

While barcode saturation establishes that background barcodes are abundant, it does not quantify how much background signal enters real cells. Human-mouse mixture datasets address this directly, as any cross-species counts within a barcode must originate outside the cell it labels. In an Evercode v1 mixture dataset (~ 7,000 cells) (Tran et al., 2022), every barcode carried cross-species counts (Fig. 7A). Restricting to barcodes with less than 10% cross-species signal, which are likely to represent singlets, median cross-species contamination was 1.2% for human barcodes (IQR 0.9–1.5%) and 2.1% for mouse barcodes (IQR 1.7–2.7%). In absolute terms, interquartile ranges were 29–57 cross-species counts for human barcodes and 38–74 for mouse barcodes. In the 90th percentile, contamination reached 80 counts for human barcodes and 102 for mouse barcodes, indicating that a subset of otherwise singlet-like barcodes carry appreciable background. An Evercode v2 mixture dataset gave comparable values (IQR 1–3%; 28–63 counts) (Fig. 7B). All of these figures are lower bounds, since intra-species contamination is undetectable by this approach and genes mapping to both genomes were excluded.

**Fig 7.**
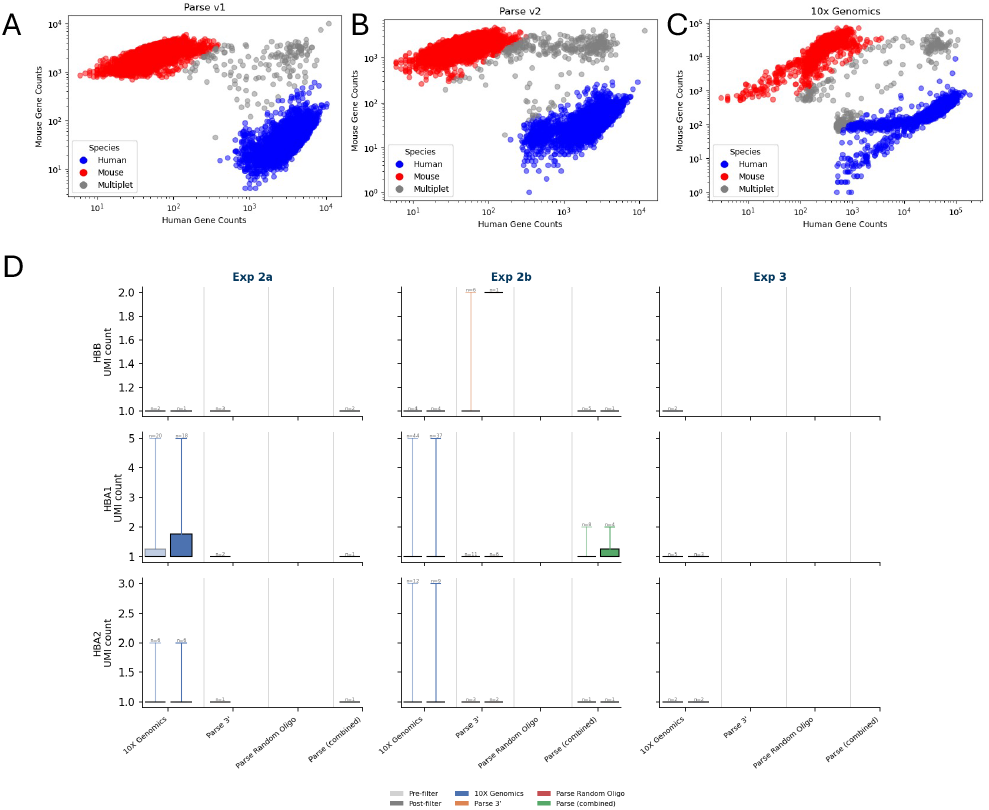
Parse exhibits substantial markers of ambient RNA contamination, inconsistent with claims of reduced contamination relative to 10X. (A) Log-log scatter plot of counts aligned to the mouse genome vs. the human genome for the Parse Evercode WT v1 human–mouse mixture dataset. Barcodes with <10% mouse counts are labelled human (blue), those with <10% human counts are labelled mouse (red), and the remainder multiplets (grey). (B, C) As in (A), for the Parse Evercode WT v2 and 10X Genomics Chromium Single Cell 3’ human–mouse mixture datasets respectively. (D) Box plots of HBB, HBA1, and HBA2 counts across the three PBMC experiments, restricted to cells with at least one count of the respective gene, before (light) and after (dark) filtering. Each box plot is labelled with the number of cells shown.

For comparison, a Chromium v3.1 mixture dataset (~ 5,000 cells) gave interquartile cross-species contamination of 0.5–0.8% for human barcodes and 1–2% for mouse barcodes, corresponding to interquartile ranges of 164–327 and 223–385 counts respectively (Fig. 7C). The higher counts at lower percentages reflect the substantially greater sequencing depth of this dataset, which makes the percentage and count scales non-comparable across platforms. The two platforms are therefore of broadly similar magnitude on this measure, and we do not read these data as establishing that either platform is cleaner than the other. What they do establish is that contamination in Parse is neither absent nor negligible: it is present in every barcode of both mixture datasets, at levels comparable to those in a droplet-based library.

Barcode-rank plots were consistent with both results. Knees were generally more pronounced in 10X than in Parse libraries (Fig. S9/S10), though sharpness varied across experiments (Macosko et al., 2015; Klein et al., 2015), suggesting that background barcodes are less cleanly separable from cells in Parse. That contamination in Parse is substantial is difficult to reconcile with reported comparisons attributing markedly lower contamination to Evercode WT. The discrepancy is explained by the metrics on which those comparisons rest. Lower contamination in Evercode WT has been attributed to lower mitochondrial and ribosomal transcript percentages (Parse Biosciences, 2026), but neither is specific to contamination. Mitochondrial percentage varied by dataset in our data with no platform trend (see above). Ribosomal percentage varies by cell type and so doesn’t make for a good contamination marker (Ilicic et al., 2016). In addition, both split-pool platforms in a broader cross-technology comparison – Parse and Scale Biosciences – showed RP percentages near 0%, against ~ 10% for microwell-, hydrogel-, and droplet-based methods (De Simone et al., 2025; Petibon et al., 2021). Low RP percentage therefore tracks the split-pool platform class, and in our data is accounted for by the nuclear-capture bias described above.

Hemoglobin transcripts avoid this confound, as mature PBMCs do not express them, and have been used to compare platforms on that basis (Parse Biosciences, 2026). Across our PBMC datasets (Experiments 2a, 2b, and 3), HBB, HBA1, and HBA2 were sparse in both platforms (Fig. 7D). In Experiments 2a, 2b, and 3, 10X carried no more than 5 counts of any of these genes per cell and fewer than 41 contaminated cells per gene, dropping further after filtering; Parse carried fewer still, with none detected in Experiment 3. While this suggests that Parse is less susceptible to ambient contamination, the overall number of contaminating counts is too small to establish an order-of-magnitude difference in ambient RNA between platforms.

## Discussion

Across seven matched comparisons spanning multiple tissues, species, and kit versions, we find that 10X and Parse differ along two largely independent axes. The first is a platform-level batch effect in which Parse appears to be systematically depleted for cytoplasmic transcripts relative to 10X—a shift we suspect to be due to relative enrichment for nuclear RNA in Parse. A second, smaller axis distinguishes the two Parse priming strategies from one another and is best explained by transcript GC content rather than length or the presence of a poly(A) tail. Alongside these compositional biases, we find that Parse libraries carry clear markers of ambient RNA contamination, at levels that appear broadly comparable to 10X and inconsistent with reports of markedly lower contamination in Evercode WT. We consider each of these findings, and their possible mechanisms, in turn.

### Library quality and transcript diversity

Parse libraries showed lower mapping rates than 10X across all experiments, regardless of priming strategy. One possible explanation for this is that random hexamer priming preferentially captures RNA from highly repetitive genomic regions such as ribosomal RNA, which is more difficult to map uniquely. This explanation is consistent with the randO-specific drop in unique mapping under STAR, but it does not fully account for the overall reduction in uniquely mapped reads across all Parse datasets relative to 10X, including Parse 3’, which shares 10X’s priming strategy. The lower mapping rate in Parse therefore appears to be only partly attributable to priming chemistry.

In addition, because a larger fraction of Parse reads originate from intronic sequence than in 10X, and a conventional transcriptome index contains only mature transcript sequences, we found during the course of our analysis that the transcriptome mapping rate is substantially lower for Parse than for 10X. For this reason, we recommend that Parse data be quantified against a reference that includes intronic sequence — either a full genome alignment or an intron-aware transcriptome index — so that intronic reads are retained rather than discarded as unmapped.

Separately, Parse (with both randO and oligo(dT) reads combined) detected more genes than 10X in every experiment, suggesting that Parse may capture a somewhat more diverse set of transcripts. However, the greater gene count did not consistently translate into higher Shannon diversity, which was mixed across experiments (Experiment 4 in particular ran counter to the general trend). Taken together, Parse likely recovers a broader range of genes in most settings, but the effect is not cleanly reflected in entropy-based measures.

### Cytoplasmic depletion in Parse and the role of fixation

The largest and most consistent difference between the platforms is the depletion of cytoplasmic transcripts in Parse. Transcripts expected to be abundant within a cell—ribosomal protein genes, mitochondrial respiration genes, and constitutively transcribed housekeeping genes more generally—were depleted in Parse relative to 10X. In the case of ribosomal protein genes, Parse fell below even the baseline of roughly 10% bulk RP content reported across other single-cell technologies (Petibon et al., 2021; De Simone et al., 2025). In contrast, Parse exhibited enrichment of regulatory proteins, including transcriptional regulators, regulators of GTPase signaling, and protein kinase activity.

To our knowledge, the two most likely explanations for this pattern are that (1) fixation followed by permeabilization results in the loss of cytoplasmic transcripts, producing the relative nuclear enrichment we observe, or that (2) fixation produces RNA–protein crosslinks that preferentially restrict access to certain transcripts. These mechanisms are not mutually exclusive, and our data do not decisively distinguish them.

A few comparison papers (De Simone et al., 2025; King et al., 2026) have suggested the first mechanism, citing evidence that fixation causes substantial loss of cytoplasmic contents. However, much of this evidence concerns alcohol-based fixation, whose mechanism differs entirely from the formaldehyde fixation used in split-pool protocols (Rosenberg et al., 2018). The literature on formaldehyde fixation is more ambiguous, and the extent of RNA loss depends heavily on the exact protocol. In the context of in-situ hybridization, Bentley Lawrence and Singer (1985) report that RNA retention is essentially unchanged after brief paraformaldehyde fixation, whereas alcohol-based methods lose 75% or more of total cellular RNA. Urieli-Shoval et al. (1992) similarly find near-total RNA loss with alcohol-based fixatives; paraformaldehyde-fixed tissue retained more, though only around 45%, comparable to fresh-frozen samples. Using Raman imaging, Hobro and Smith (2017) likewise report that fixation degrades biochemical information most severely in the cytoplasm and that organic solvents are far more destructive than aldehyde-based methods, with aldehyde-fixed nuclei scoring close to live cells—an indication that nuclear content is comparatively well preserved.

This body of evidence establishes that formaldehyde-associated RNA loss is partial rather than total, and that nuclear content is comparatively preserved, both of which are consistent with a loss that skews cytoplasmic and would produce the nuclear enrichment we observe. We are not, however, aware of direct evidence that formaldehyde-associated RNA loss is *preferentially* cytoplasmic in the way our mechanism requires. In addition, loss of cytoplasmic transcripts during scRNA-seq is associated with an increase in detected mitochondrial transcripts, which we did not find to be enriched in the Parse datasets.

The second mechanism remains equally open. Formaldehyde is well established to create crosslinks between nucleotides and proteins, forming most readily between lysine and guanine, with crosslinking affinity in live cells governed by molecular proximity (Hoffman et al., 2015). Such crosslinks could plausibly restrict reverse-transcriptase access to particular transcripts without any loss of the RNA itself. Notably, Hobro and Smith (2017) report that permeabilization with Triton—as used in split-pool protocols—can reverse adduct formation, which complicates any simple account and suggests the balance between crosslink formation and reversal may itself depend on protocol details. Altogether, while fixation-associated cytoplasmic loss is plausible and partially supported, it is unlikely to be total, and crosslink-mediated restriction of transcript access remains a viable alternative. Both mechanisms merit further investigation.

### Priming-strategy bias and GC content

The bias between oligo(dT) and random hexamer priming within Parse is orthogonal to the platform-level bias between 10X and Parse. Contrary to expectation, this priming bias does not appear to be driven by transcript length, by the presence or absence of a poly(A) tail, or by the adenine-rich regions responsible for internal off-target priming by oligo(dT) primers. Instead, it tracks most directly with GC content. Two mechanisms could account for this: random hexamers may form more stable duplexes with GC-rich sequence, or the secondary structure of GC-rich transcripts may inhibit full-length reverse transcription initiated from the poly(A) tail.

This is notable given we also observed that different Parse wells were sequenced to differing depths and with differing proportions of oligo(dT)- and random-hexamer-primed counts (Fig. S12). This raises the concern that sample multiplexing may introduce systematic biases—including in GC content—that reflect the ratio of the two priming strategies across wells rather than genuine biological differences.

It is also notable that the two orthogonal biases converge on the same gene classes. Just as 10X is enriched for ribosomal and mitochondrial respiration genes relative to Parse, oligo(dT) priming is enriched for these same classes relative to random hexamer priming, independent of GC content. The most likely explanation is that these transcripts are highly expressed across cell types and therefore make otherwise subtle biases more visible, rather than that the two biases share a mechanism.

### Depletion of canonical cell-type markers

A further consequence of the platform-level bias is that 10X was significantly enriched for canonical PBMC marker genes relative to Parse. This trend has been noted elsewhere: Xie et al. (2024) found that while Parse was better able to detect lowly expressed genes and resolve rare cell types than 10X, it was notably depleted for CD3D, CD3G, CD3E, and NKG7—markers conventionally used to identify T cells and natural killer cells—and Filippov et al. (2024) similarly reported depletion of the CD3 genes in Parse relative to 10X. The convergence of three independent comparisons on the same result suggests this is a robust property of the platforms rather than a dataset-specific effect. Its practical implication is that, although both technologies provide sufficient information to resolve distinct cell types, cell atlases and cell-type annotations derived from 10X and Parse are not directly comparable.

### Ambient RNA contamination

Parse exhibits clear signs of RNA contamination, most likely of magnitude comparable to 10X. One comparison paper (Filippov et al., 2024) attributed contamination in split-pool data to index hopping rather than conventional ambient contamination. We find this unlikely to be the primary mechanism. To our knowledge there is no evidence that index hopping is a significant source of contamination in Parse, or that it is more prevalent than ambient contamination; moreover, index hopping affects only the final-round Illumina sample index, and so cannot account for the near-complete saturation of barcode space we observe. Background contamination was also present in experiments using a single sublibrary, where index hopping between sublibraries cannot occur. Most tellingly, Rebboah et al. (2026) reported significant cross-tissue contamination in a tissue-multiplexing experiment, which points to contamination arising upstream of library generation rather than during sequencing. The mechanism of contamination in Parse thus remains uncharacterized, but the available evidence points away from index hopping and toward a process that acts before or during barcoding. Given that Parse exhibits reduced contamination from hemoglobin transcripts—which arise from ambient RNA contamination introduced during preparation of the cell suspension—it is possible this noise is primarily due to a downstream source uncommon to droplet-based technologies, such as barcode leakage between wells.

### Additional considerations

Although a key measure of dataset quality, we did not attempt to compare the multiplet rate between the two technologies, as computational methods are not designed to reliably estimate the number of multiplets a technology produces. Human–mouse mixture experiments remain the gold standard for this purpose. The mixture datasets we examined (Tran et al., 2022), which recovered roughly 5,000–7,000 cells per species, imply cross-species multiplet rates below 2% for the v1 and v2 chemistries and thus true multiplet rates below roughly 4%, though these figures vary with the number of cells loaded. For comparison, Zheng et al. (2017) reported an expected multiplet rate of around 8% at a similar cell count, albeit for an earlier generation of the droplet technology underlying 10X. This is consistent with the expectation that multiplets in Parse arise only through barcode collisions, whereas in droplet-based platforms they can arise through both collision and co-encapsulation. One dataset was a striking exception: the human–mouse mixture in the 10X/Parse comparison of Hornung et al. (2023) reported a Parse multiplet rate of 49%. Applying a cross-species count threshold in our own reanalysis, we recovered a comparably high figure of roughly 30%, against roughly 6% in the matching 10X dataset (Fig. S11). We applied the same threshold to the other mixture datasets we examined without obtaining anything approaching this rate, and we are not aware of another study reporting a comparable value for Parse. We are unable to account for why this is the case.

In addition, we did not compare the number of cells recovered or the number of UMIs per cell across platforms, despite these being common points of comparison. The number of cells recovered is highly dependent on the number of cells loaded, which was not consistently reported or directly comparable across the source studies. The number of recovered cells also depend heavily on the filtering threshold used to separate cells from background. As shown in Figs. S9 and S10, the knee is often less clearly defined in Parse than in 10X, making it more difficult to choose an appropriate threshold. Per-cell UMI yield is likewise not directly comparable here, as each dataset was subsampled to a common read depth within its experiment, so per-cell counts might be more reflective of the artificial library size than capture efficiency.

Finally, several of the datasets analyzed here multiplexed samples, using hashtag oligonucleotides in 10X and well-based multiplexing in Parse. We did not attempt to demultiplex samples corresponding to distinct biological replicates or perturbations, as a comparison of multiplexing strategies was beyond the scope of this work and hashtag-based and well-based multiplexing are not directly comparable. This may have introduced additional confounding.

## Conclusion

To our knowledge, this is the first comparison of droplet-based and split-pool platforms to separate reads by priming strategy, allowing platform-level and priming-level biases to be distinguished directly. We find that the two axes are largely independent, and that the platform-level difference is by far the larger of the two. While priming strategy introduces a GC-content bias within Parse, the dominant difference between the technologies is a systematic compositional shift that holds across priming strategies. This shift is characterized by the depletion in Parse, relative to 10X, of ribosomal protein and mitochondrial respiration genes as well as for other ubiquitous protein-complex transcripts in conjunction with elevated intronic content and MALAT1 expression. We take this to indicate that Parse preferentially targets nuclear rather than cytoplasmic RNA, likely due to depletion of cytoplasmic RNA. As there is previous literature on fixatives resulting in perturbation of cytoplasmic contents, we hypothesize that this could be due to loss of RNA from the cytoplasm. However, as the aldehyde fixatives used in the Parse protocol are less susceptible to this kind of RNA loss, it could instead be the case that aldehyde crosslinks prevent access to certain transcripts. Finally, we assessed the claim that Parse captures markedly less ambient RNA than 10X, and do not find it supported. By measures that do not assume a droplet-specific contamination model, we find contamination in Parse is of a magnitude comparable to 10X, though its mechanism remains uncharacterized. Taken together, these results indicate that while both technologies resolve cell types effectively, the compositional differences between them are substantial enough that expression measurements and cell-type composition are not directly interchangeable across platforms.

## Methods

### Read retrieval and library assembly

SRA runs were downloaded with prefetch and converted to FASTQ with fasterq-dump (`--split-files --skip-technical`; SRA-Tools v3.0.3 Leinonen et al. (2010)) and compressed with pigz. ENA runs were streamed directly from the ENA FTP server as gzipped FASTQ. For assays distributed as multiple sublibraries, all sublibraries belonging to a single assay were combined into one paired-end FASTQ with `splitcode --remultiplex` (splitcode v0.30.0 Sullivan and Pachter (2024)), preserving the original read-1/read-2 structure. Because a Parse sublibrary is resolved by its sequencing index rather than by the combinatorial barcodes themselves, identical round-1/2/3 barcode combinations occurring in different sublibraries label different cells. Parse sublibraries were therefore combined with `--bclen=4`, which prepends a distinct 4 bp sequence to read 2 of every sublibrary; runs originating from the same sublibrary were assigned the same sequence. This synthetic barcode was carried through the remainder of the pipeline as a fourth cell barcode.

### Reference genomes and indices

Reference genome FASTA and GTF annotations were obtained from Ensembl Harrison et al. (2024). Human assays used GRCh38 (release 115) and the mouse thymocyte assay used GRCm39 (release 115). For the barnyard analysis a combined reference was constructed by concatenating the human (GRCh38, release 112) and mouse(GRCm39, release 115) genomes; every chromosome name, `gene_id`, and `gene_name` was prefixed with a species tag (`HUMAN_ / MOUSE_`) so that reads could be assigned unambiguously to a species of origin.

For pseudoalignment, a nascent-and-mature (nac) index was built for each reference with kb ref `--workflow nac` (kb-python v0.29.3 Sullivan et al. (2025), wrapping kallisto v0.51.1 Bray et al. (2016) and bustools v0.44.1 Melsted et al. (2021)). The nac workflow indexes both spliced (mature) and unspliced (nascent) transcript sequences, allowing intronic reads to be quantified separately from exonic reads.

### Parse split-pool barcode handling

Parse Evercode encodes cell identity through three rounds of combinatorial barcoding carried on read 2, with the round-1 (RT) barcode existing in two chemistries within each well: an anchored oligo(dT) primer (polyT) and a random-hexamer primer (randO). Because the polyT and randO primers in a given well label the same cell, they must be reconciled during quantification.

Barcode whitelists and the kallisto technology string for each kit/chemistry were generated programmatically from the Parse Biosciences Evercode WT barcode reference found in the Parse Biosciences software (version 1.7.2), keyed on the kit name and, where applicable, the subset of sample wells used. All read-2 barcode and UMI coordinates were offset by 4 bp to account for the sublibrary barcode prepended during library assembly, and the sublibrary barcode itself was registered as an additional cell barcode occupying the first 4 bp of read 2. Reads were then processed with `splitcode` (v0.30.0) in two steps: (i) reads were filtered to those carrying an expected round-1 barcode at the kit-specific position, allowing one mismatch (Hamming distance *≤* 1); and (ii) the retained reads were partitioned into polyT and randO subsets for separate downstream analysis.

Filtered Parse reads were quantified with kb count `--workflow nac` using the kit-specific technology string, forward strandedness (`--strand=forward`), and single-end parity (`--parity=single`). A per-round barcode on-list (`-w onlist.txt`) was supplied for barcode error correction, listing the sublibrary barcode alongside the three combinatorial rounds so that cells were resolved by sublibrary as well as by barcode combination. A replacement map (`-r replace.txt`) collapsed each randO round-1 barcode onto its polyT counterpart in the same well so that both primer chemistries contributed counts to a single cell. Three count matrices were produced per Parse assay: the full filtered set, the polyT-only subset, and the randO-only subset.

### 10x Genomics quantification

10x reads were quantified with `kb count --workflow` nac using the appropriate technology string (10XV3 for Chromium Next GEM v3 assays, 10XV4 for the Chromium v4 assay) and the matching cell-barcode on-list. As with the Parse data, the nac workflow yielded mature, nascent, and ambiguous count layers.

### Read-depth matching

To compare technologies at equal sequencing effort, processed read counts were tabulated for every assay in an analysis (the full filtered FASTQ for each 10x and Parse assay, plus the Parse polyT and randO subsets) and the minimum across assays was taken as the target depth. Each FASTQ was then downsampled to this common depth with `seqtk sample` using a fixed random seed (42) and re-quantified with kb-python as above. The 10x, full-Parse, polyT, and randO read sets were each subsampled to the same depth so that all pergene and per-cell comparisons reflect differences in chemistry rather than in coverage.

### Cell calling and quality control

Downstream analysis was performed in Python with Scanpy (v1.11.4 Wolf et al. (2018)) and AnnData (v0.12.2 Virshup et al. (2024)). Each kb-python h5ad matrix was loaded, Ensembl gene IDs were mapped to gene symbols, and genes/cells with zero counts were removed. Percell and per-gene summary statistics were computed, including total UMI counts, genes detected, and the fraction of counts attributable to nascent (unspliced), mature (spliced), and ambiguous molecules. Alignment statistics were drawn from kb-python’s `run_info.json` and, where STAR alignments were generated, from STARsolo’s `Log.final.out`.

Cells were called from each assay’s barcode-rank (knee) plot (See Figs. S8 and S9): the UMI distribution was ranked and a manually chosen assay-specific minimum-UMI threshold was applied to separate cells from empty droplets/barcodes, retaining all barcodes above the threshold.

Gene-level annotations were assembled once per reference. Gene biotype (protein coding, lncRNA, mitochondrial, ribosomal, and OXPHOS/Proteasome/Spliceosome classes, identified by biotype and gene-symbol prefixes) was parsed from the Ensembl GTF, while transcript length and GC content were computed directly from the kallisto cDNA FASTA and averaged across isoforms per gene—guaranteeing complete coverage for every gene in the index. Per-cell biotype fractions were added from these annotations.

### Gene-body coverage

To assess 5^*′*^/3^*′*^ coverage bias, depth-matched reads were aligned to the genome with STARsolo (STAR v2.7.11b Dobin et al. (2013)). 10x reads were aligned in `CB_UMI_Simple` mode with the 10x cell-barcode whitelist; Parse reads were aligned in `CB_UMI_Complex` mode with the three per-round Parse whitelists and barcode/UMI positions derived from the kit technology string (1-mismatch barcode matching). Parse alignments were generated separately for the full, polyT, and randO read subsets. Coordinate-sorted BAM files were indexed with SAMtools (v1.21 Danecek et al. (2021)). A BED12 model was produced from the GTF with `gffread` (Pertea and Pertea, 2020) and subset to housekeeping transcripts from the HRT Atlas (Hounkpe et al., 2021); transcript-body coverage was then computed across all assays with RSeQC `geneBody_coverage.py` (RSeQC v5.0.4 Wang et al. (2012)).

**Pergene Composition Comparison**. Pergene expres-sion was summarised as log-counts-per-million (log-CPM): for each gene *g*, ln (CPM_g_ + 1) where CPM =(Σ_cells_ UMI_*g*_)/(Σ^all^ UMI) × 10^6^. Log-CPM vectors from the two technologies were aligned on shared gene symbols (genes absent from one assay assigned a value of zero), and Lin’s concordance correlation coefficient (Lin, 1989) was computed:

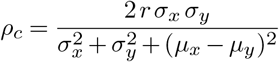

where *r* is the Pearson correlation and *σ, μ* denote the standard deviation and mean of each log-CPM vector. The CCC measures both correlation and agreement along the line of perfect concordance (*y* = *x*), penalising systematic differences in level or scale that *r* alone would not detect.

### Cross-study differential composition and functional enrichment

To identify genes that were consistently over- or under-represented by one technology across studies, a pseudobulk differential-expression analysis was performed with edgePython (Pachter, 2026), a Python implementation of the edgePython workflow (Robinson et al., 2010; Lun et al., 2016). Counts were summed across all cells within each assay to form one pseudobulk profile per sample, and profiles were restricted to genes detected in every sample. A DGE-List was constructed and normalised by the trimmed mean of M-values (TMM) method (Robinson and Oshlack, 2010). A design matrix of the form ~ `tissue + technology` was used so that the technology effect was estimated while controlling for biological source; lowly expressed genes were removed with `filterByExpr`. Negative-binomial dispersions were estimated (`estimateDisp`), a quasi-likelihood negative-binomial GLM was fit (`glmQLFit`), and the technology coefficient was tested with the quasi-likelihood F-test (`glmQLFTest`). P-values were adjusted by the Benjamini– Hochberg procedure, and genes with FDR below the chosen threshold were called as differentially represented between technologies.

Genes consistently enriched in each technology were then tested for functional enrichment against the GO Biological Process, GO Molecular Function and GO Cellular Component gene-set libraries, retrieved with gseapy (Fang et al., 2023) from Enrichr (Kuleshov et al., 2016). Because GC content is itself predictive of differential representation in these comparisons, a conventional hypergeometric test would report terms that are enriched merely because their genes are GC-extreme. Enrichment was therefore computed with goseq (Young et al., 2010), which fits a monotonic probability weighting function *P* (DE | GC) over the tested genes and evaluates each term with a Wallenius non-central hypergeometric test whose odds are the ratio of mean weights inside and outside the term. Median-transcript GC content was used as the bias variable, and all genes retained by `filterByExpr` formed the statistical universe. The uncorrected hypergeometric test was computed on the same universe and gene sets as an uncorrected reference. P-values were adjusted by the Benjamini–Hochberg procedure within each ontology, and terms were classified as GC-robust (adjusted p below 0.05 both before and after correction, i.e. enrichment orthogonal to the GC bias) or GC-driven (significant only before correction).

Because Gene Ontology output is highly redundant— nested parent and child terms, and semantically unrelated terms annotated to the same genes, are reported as separate results—significant terms were grouped by the overlap coefficient of their differentially expressed gene sets, |*A ∩ B*| */* min(|*A*|, |*B*|) *≥* 0.7. Seed groups were taken as the maximal cliques of the resulting similarity graph, so that every pair of terms within a seed exceeds the threshold; seeds whose combined gene sets overlapped by more than 40% (Jaccard) were then merged iteratively, following the multiple-linkage merging of DAVID’s fuzzy functional classification (Huang et al., 2009, 2007). Without this merging step, clique enumeration on a dense similarity graph returns many near-duplicate groups spanning essentially the same genes. Groups sharing the same representative term were additionally folded together, so that each term represents at most one group. A term may otherwise belong to more than one final group. The most significant term in each group is reported as its representative, together with the genes present in at least half of the group’s member terms; the strict intersection over all members is typically empty once seeds have been merged. The overlap coefficient was used in preference to the Jaccard index because Gene Ontology redundancy is dominated by nested terms, for which the Jaccard index penalises the difference in set size, and in preference to semantic-similarity approaches because these terms co-occur through shared genes rather than through biological relatedness.

#### Cell-typing of Human PBMC Datasets

For the human PBMC datasets (Exp 2a, Exp 2b, and Exp 3), cells were annotated with CellTypist (v1.7.1 Xu et al. (2023); Conde et al. (2022)) using the Immune_All_High model. Annotation was performed after cell filtering, normalization to 10^4^ counts, and log1p transformation.

## Data Availability

This study analyzed only previously published, publicly available single-cell RNA-seq datasets; no new sequencing data were generated. Raw sequencing reads were obtained from the NCBI Sequence Read Archive (SRA) and the European Nucleotide Archive (ENA) under the following accessions:

- **Exp 1** (Reh B-ALL cell line, *ex vivo* drug perturbation; 10x Chromium Next GEM 3^*′*^ v3 with MULTI-seq and Parse Evercode WT Mini v1): GEO GSE229617 (BioProject PRJNA955181), runs SRR24154339– SRR24154342 Gezelius et al. (2024).
- **Exp 2a/2b** (cryopreserved human PBMCs from two donors; 10x Chromium Next GEM 3^*′*^ v3 and Parse Evercode WT v2): GEO GSE246624 (BioProject PRJNA1033995), runs SRR26594138– SRR26594155 Xie et al. (2024).
- **Exp 3** (cryopreserved human PBMCs; 10x Chromium Next GEM 3^*′*^ v3.1 and Parse Evercode WT v2): BioProject PRJNA1106903 (SRA study SRP505235), runs SRR28867563 (10x) and SRR28867558 (Parse) De Simone et al. (2025).
- **Exp 4a/4b** (mouse thymocytes; 10x Chromium Next GEM 3^*′*^ v3 with TotalSeq-B hashtags, Parse Evercode WT v2 and WT Mini v2): GEO GSE253406 (BioProject PRJNA1065472), runs SRR27585864– SRR27585896 Filippov et al. (2024).
- **Exp 5** (K562 and mESC human–mouse mixture; 10x Chromium Next GEM 3^*′*^ v3 and Parse Evercode WT Mini v2): ENA PRJEB67544 (10x; run ERR12398015) and PRJEB67549 (Parse; runs ERR12167397 and ERR12167398) Hornung et al. (2023).

## Code Availability

All code needed to reproduce the processed count matrices, analyses and figures from these accessions is available at https://github.com/mcaskey6/10XvParse.

## Author contribution

Conceptualization, I.H. Analysis, M.C. Project administration and supervision, I.H. Writing—original draft, M.C.— review and editing, M.C. and I.H.

## Acknowledgements

We thank the Pachter Lab for helpful feedback on this project. Thanks to the Caltech Bioinformatics Resource Center and the Impact of Genomic Variation on Function (IGVF) Consortium for financial and resource support for this project.

## Ethics approval and consent to participate

Not applicable. This study used only publicly available, de-identified datasets; no new experiments involving human participants or animals were performed by the authors.

