## Supplementary Figures for "A Unified Comparison of 10X Genomics and Parse Biosciences scRNA-seq Technologies"

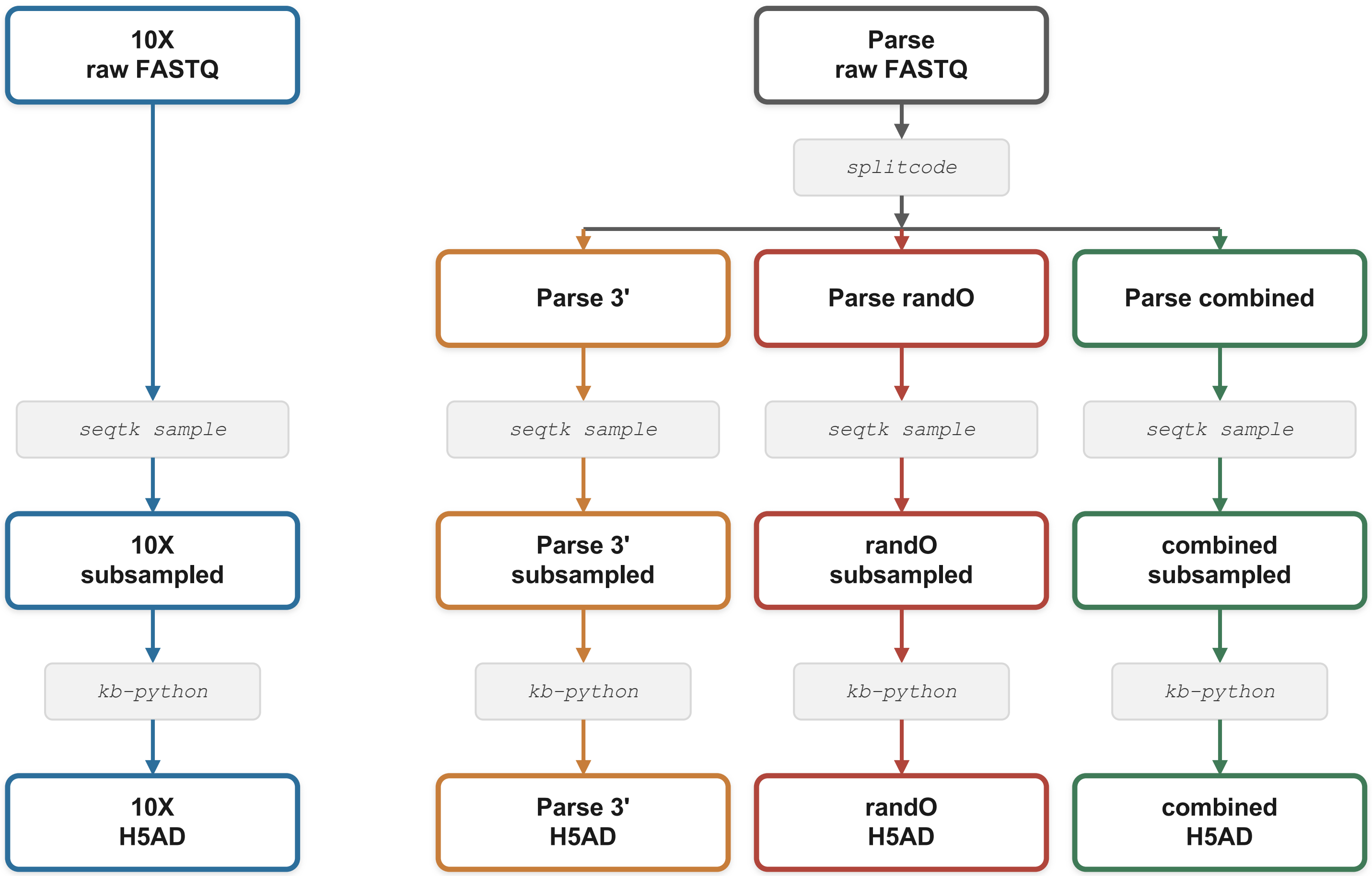


**Fig S1.** Diagram of count matrix generation for a single dataset comparison. All datasets are subsampled to the minimum read depth of the 10X, Parse 3', Parse randO, and Parse combined datasets.

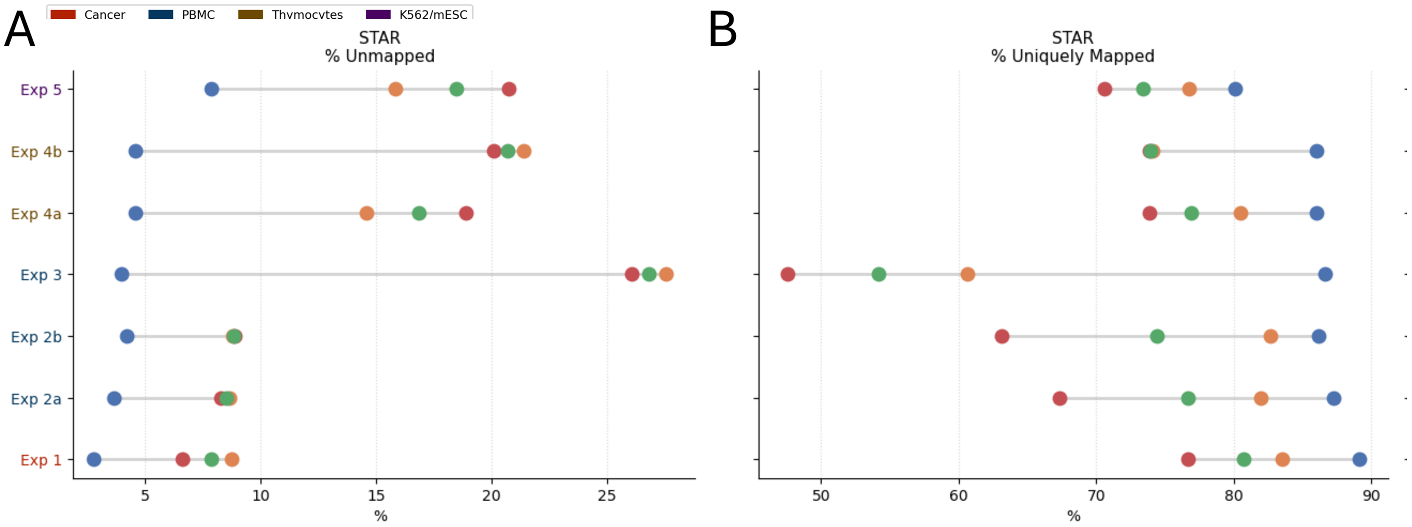


**Fig S2.** Dot plots showing the range of each STAR alignment metric across the subsampled datasets from each experiment, with experiment labels colored by tissue: (A) percent of reads unmapped; (B) percent of uniquely mapped reads

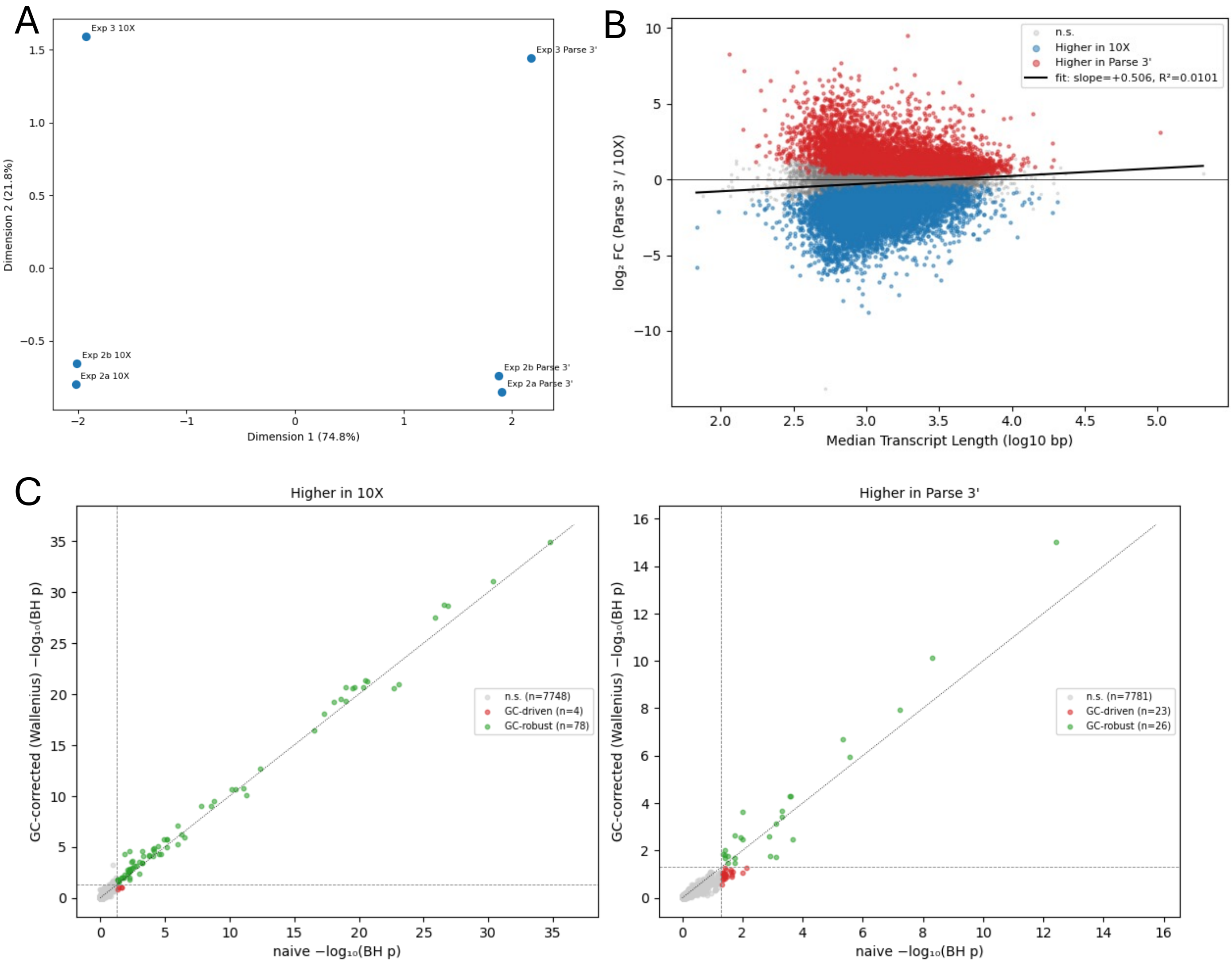


**Fig S3.** Bulk edgeR analysis comparing 10X to Parse 3' across the three human PBMC experiments, blocking on study of origin. (A) MDS Plot. (B) Median transcript length against log-fold change (Parse 3'/10X), with line of best fit. significant genes enriched in 10X are labeled in blue and enriched in Parse labelled in red. (C) For the goseq GO term enrichment analysis, comparison of $\log_{10}$ adjusted p-values (Benjamini-Hochberg) from the GC-corrected Wallenius noncentral hypergeometric test against the naive hypergeometric test, for both genes enriched in 10X and genes enriched in Parse.

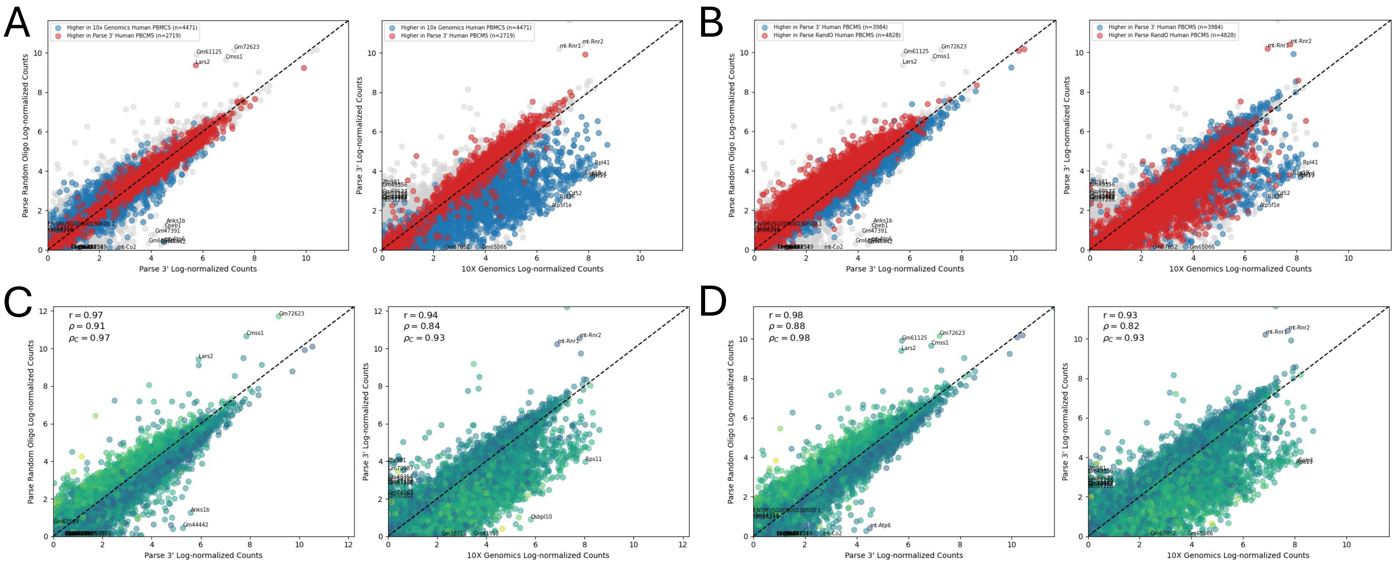


**Fig S4.** On the Exp 5a and 5b mouse thymocyte datasets: filtered logCPM bulk gene counts compared between Parse randO and Parse 3' and Parse 3' and 10X. (A) For Exp 5b, with genes colored by whether their ortholog was significantly differentially expressed (FDR ¡ 0.01) in the 10X vs. Parse 3' edgePy analysis of the four human PBMC experiments, in favor of 10X (blue) or Parse 3' (red); orthologs lacking a match or not significantly differentially expressed are shown in grey. (B) Same as (A), but with orthologs colored by the Parse 3' vs. Parse randO edgePy analysis. (C) For Exp 5a, with genes colored by median transcript GC content. (D) Same as (C), but for Exp 5b.

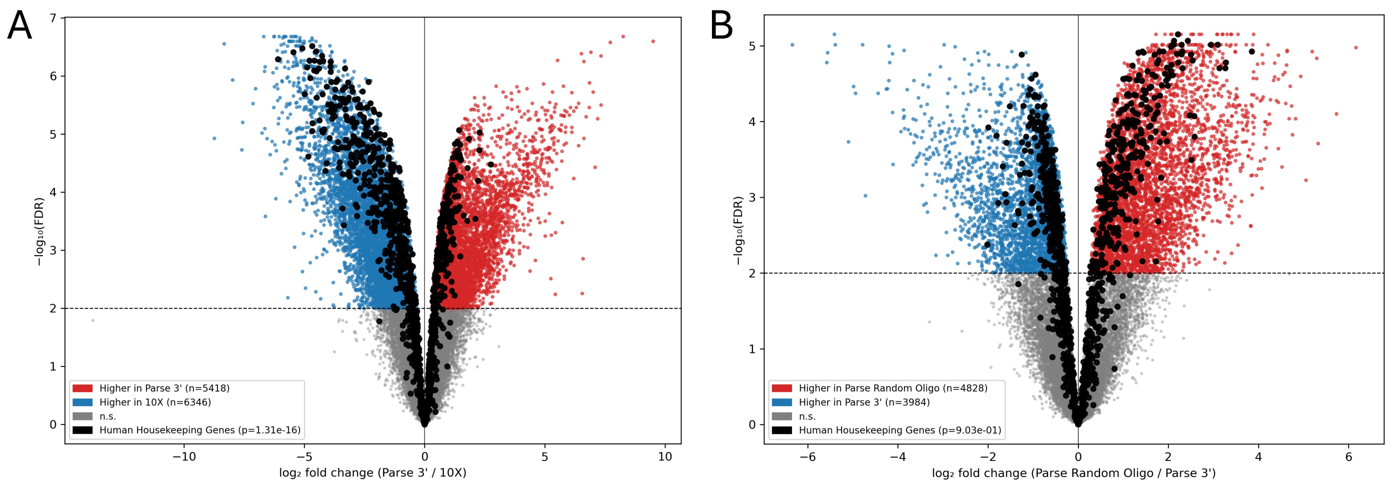


**Fig S5.** (A) From the bulk edgePy analysis of 10X vs. Parse 3' across the four human PBMC experiments, blocking on study of origin: volcano plot at an FDR threshold of 0.01, with significant genes enriched in 10X labelled in blue, those enriched in Parse in red, and housekeeping genes labelled in black. (B) Same as (A), but for Parse 3' vs. Parse randO.

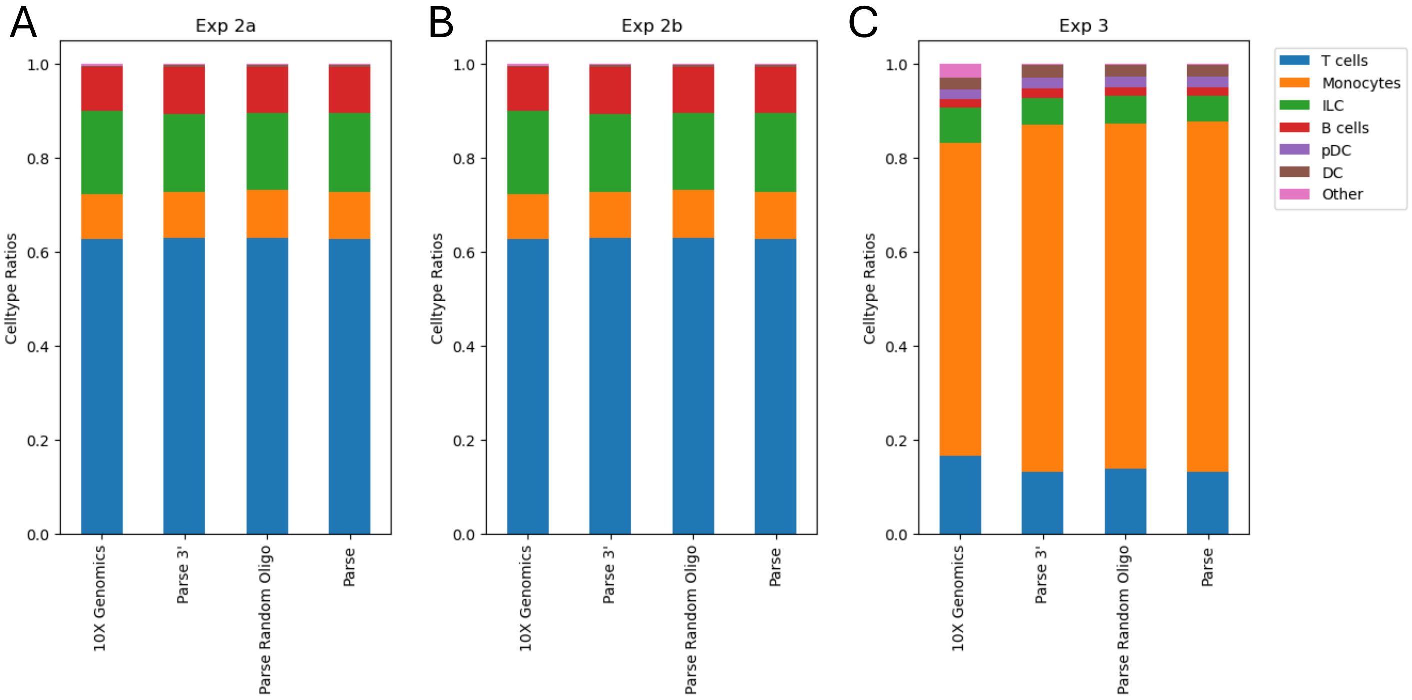


**Fig S6.** Celltype proportions for the three human PBMC comparisons based on CellTypist majority voting clustering. (A) Proportions for Experiment 2a across all subsampled datasets. (B) Same as (A) but for Experiment 2b. (C) Same as (A) but for Experiment 3.

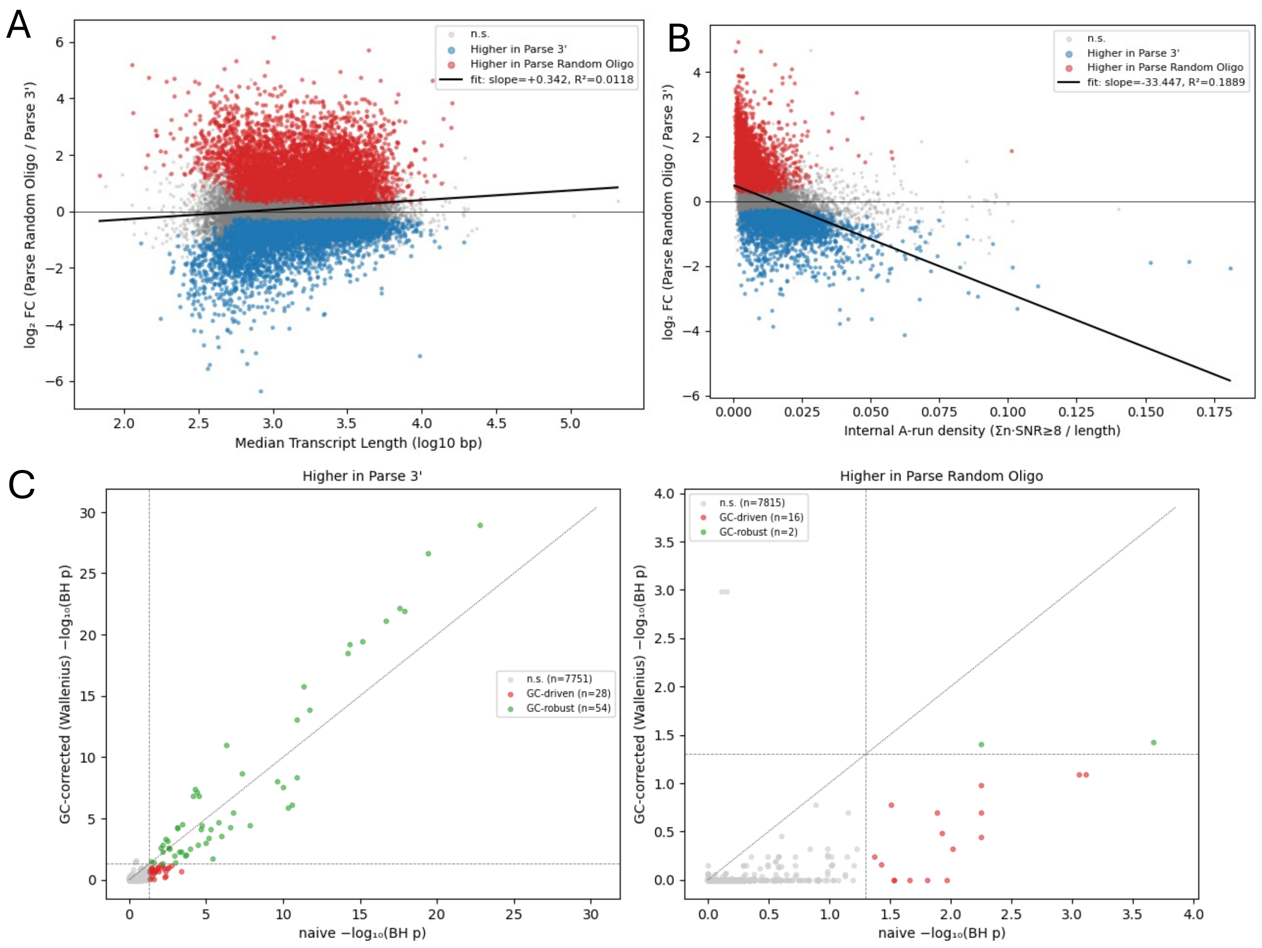


**Fig S7.** (A) Median transcript length against log-fold change (Parse randO/Parse 3'), with line of best fit. Significant genes enriched in Parse 3' are labelled in blue and those enriched in Parse randO in red. (B) Density of internal poly(A) repeats (greater than 8 nucleotides) against log-fold change (Parse randO/Parse 3'), with line of best fit. Significant genes enriched in Parse 3' are labelled in blue and those enriched in Parse randO in red. (C) For the goseq GO term enrichment analysis, comparison of $\log_{10}$ adjusted p-values (Benjamini-Hochberg) from the GC-corrected Wallenius noncentral hypergeometric test against the naive hypergeometric test, for both genes enriched in Parse 3' and genes enriched in Parse randO.

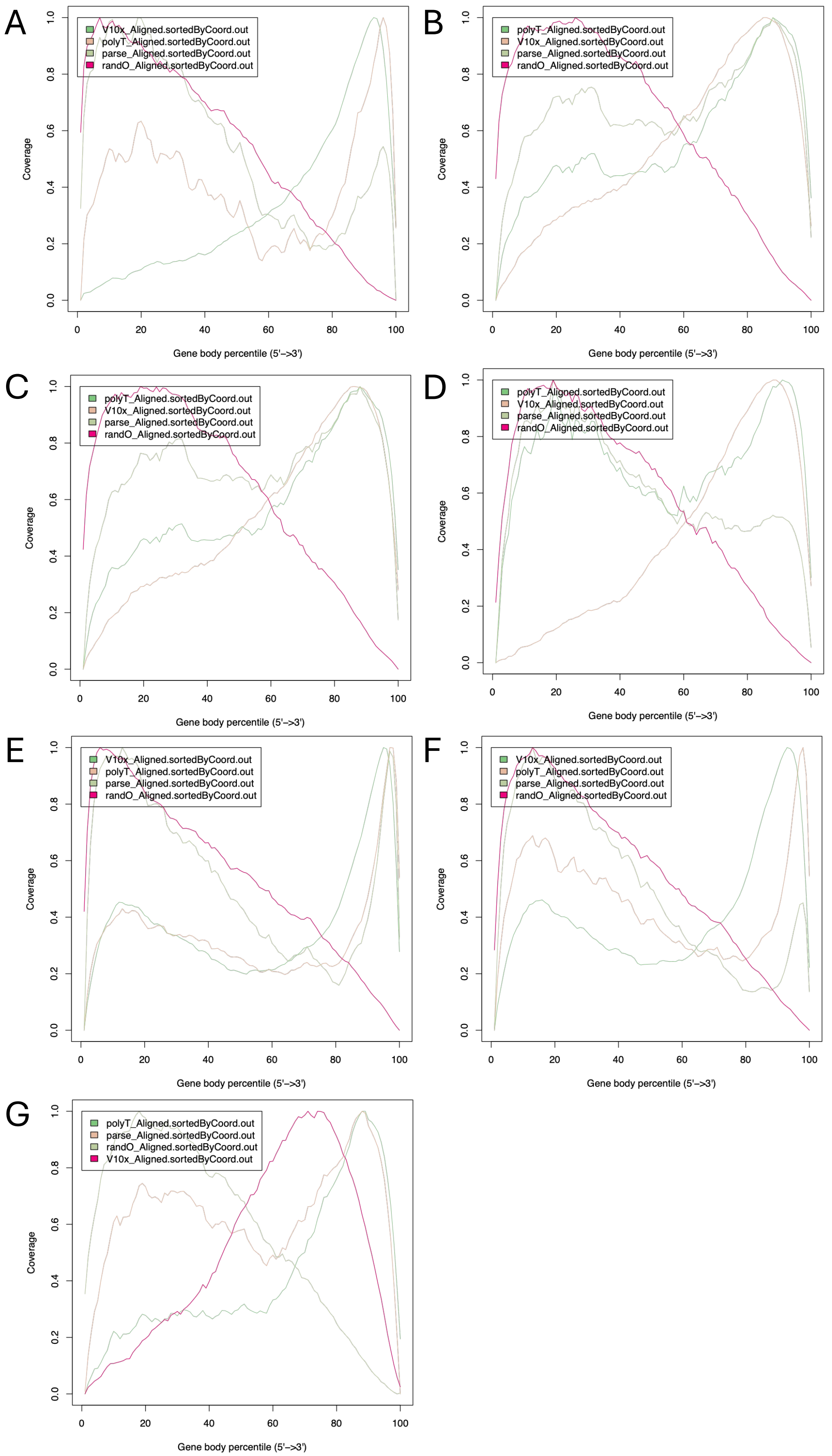


**Fig S8.** (A) Gene body coverage distributions for each dataset of Exp 1 (V10x = 10X, parse = Parse combined, polyT = Parse 3', randO = Parse randO). (B–H) Same as (A), but for Exp 2a, Exp 2b, Exp 3, Exp 4, Exp 5a, Exp 5b, and Exp 6 respectively.

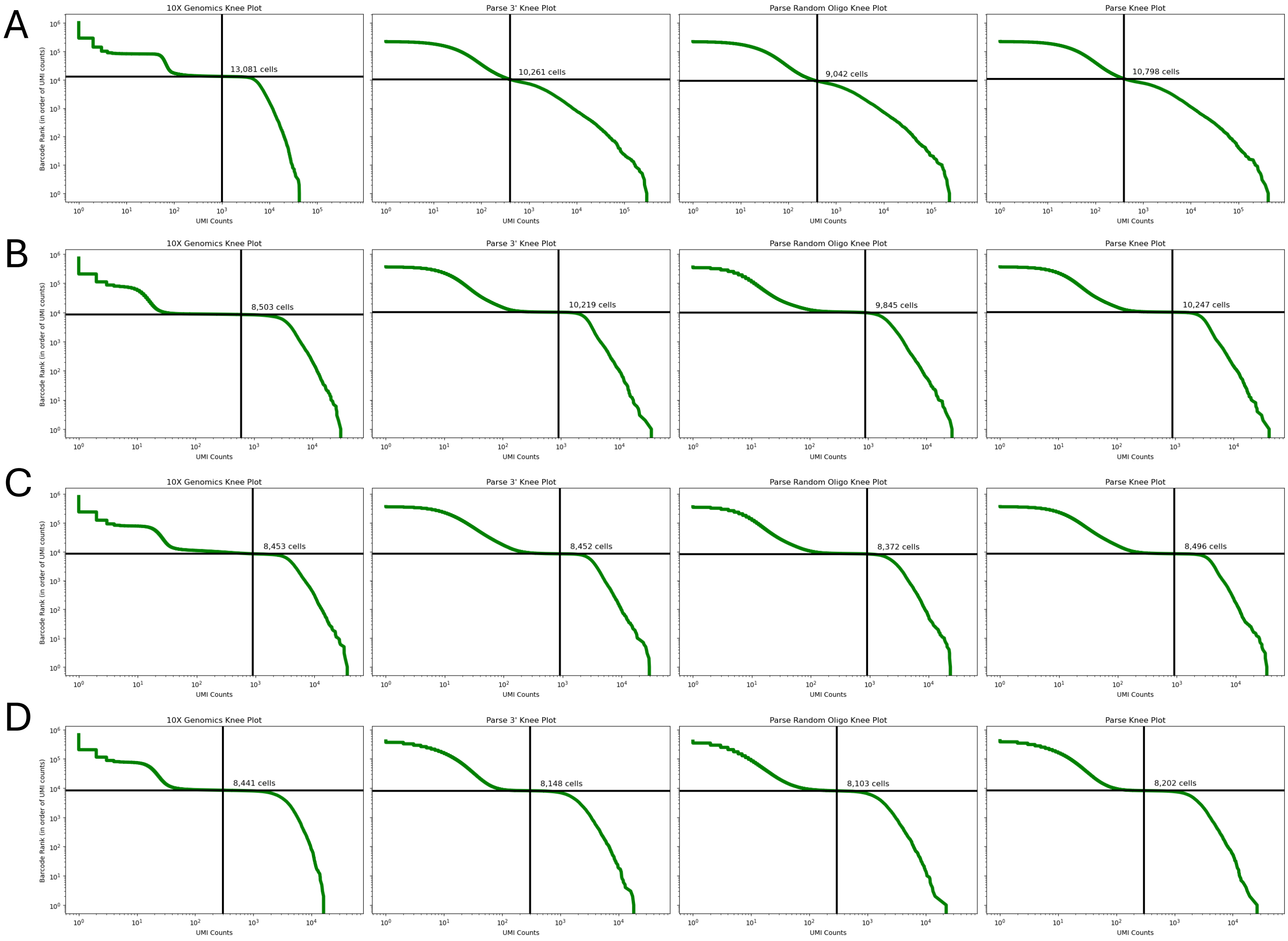


**Fig S9.** (A) Knee plots for each of the four datasets (10X, Parse 3', Parse randO, Parse combined) of Exp 1, with the filtering threshold marked in black. (B–D) Same as (A), but for Exp 2a, Exp 2b, and Exp 3 respectively.

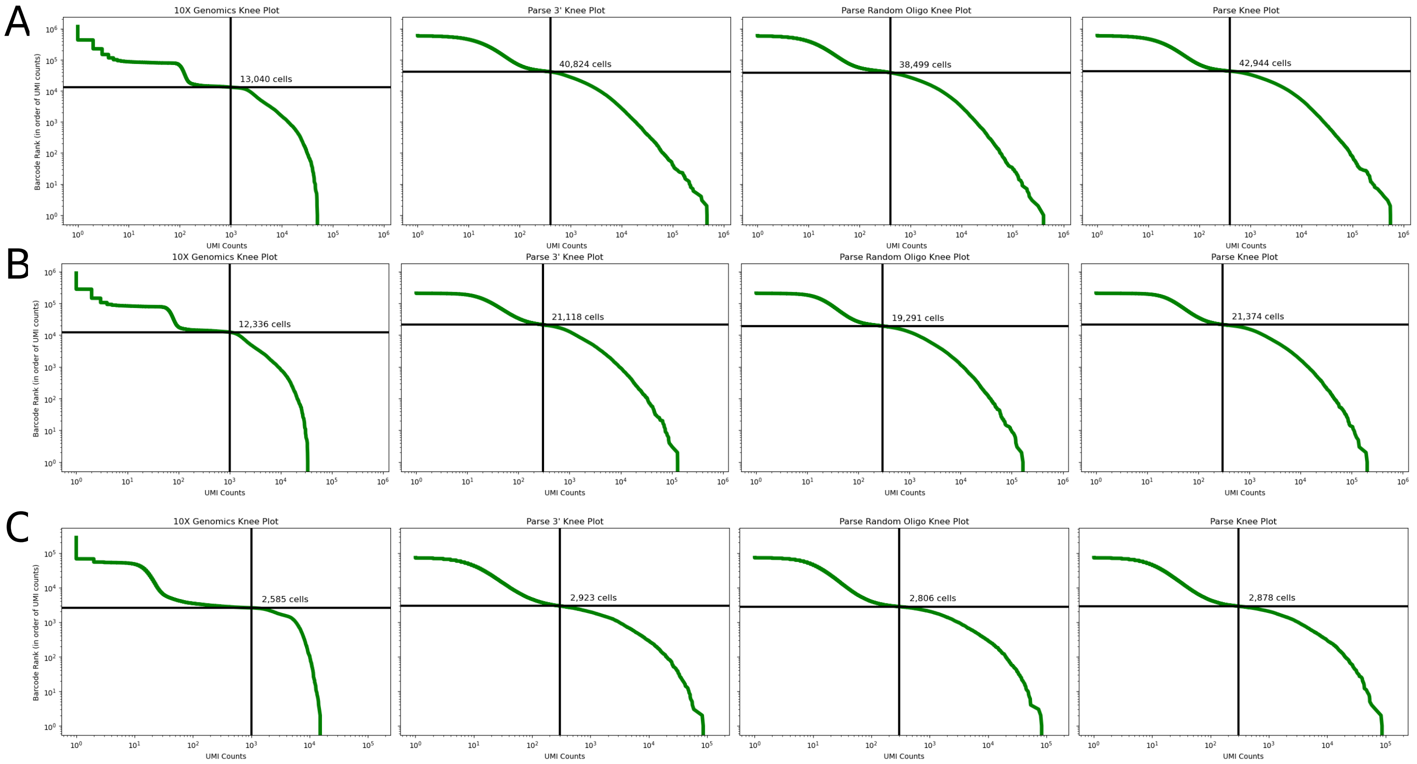


**Fig S10.** (A) Knee plots for each of the four datasets (10X, Parse 3', Parse randO, Parse combined) of Exp 4, with the filtering threshold marked in black. (B–D) Same as (A), but for Exp 5a, Exp 5b, and Exp 6 respectively.

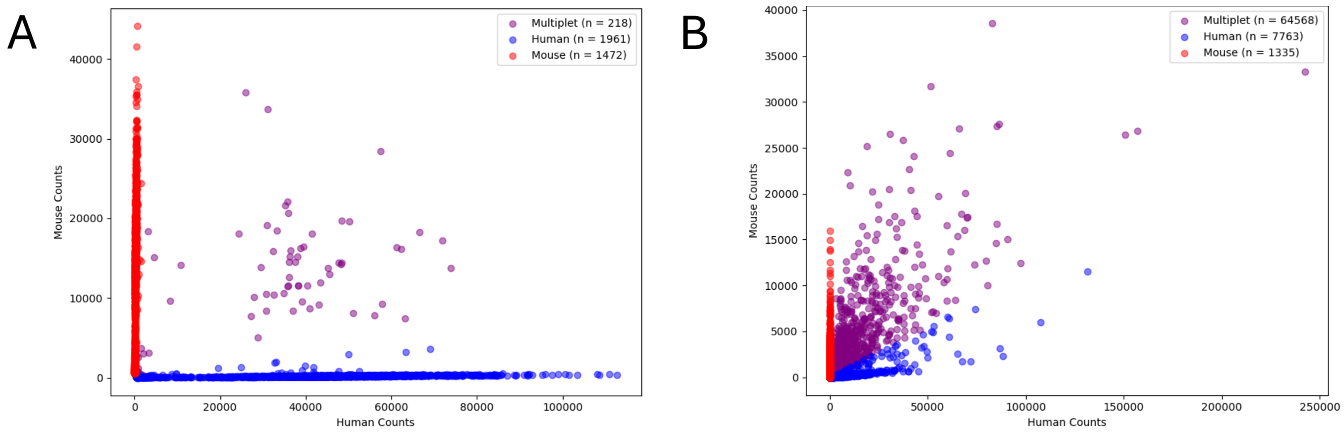


**Fig S11.** (A) Barnyard plot of the Exp 6 10X dataset; barcodes with greater than 10% off-species counts are labelled as multiplets. (B) Same as (A), but for the Parse dataset.

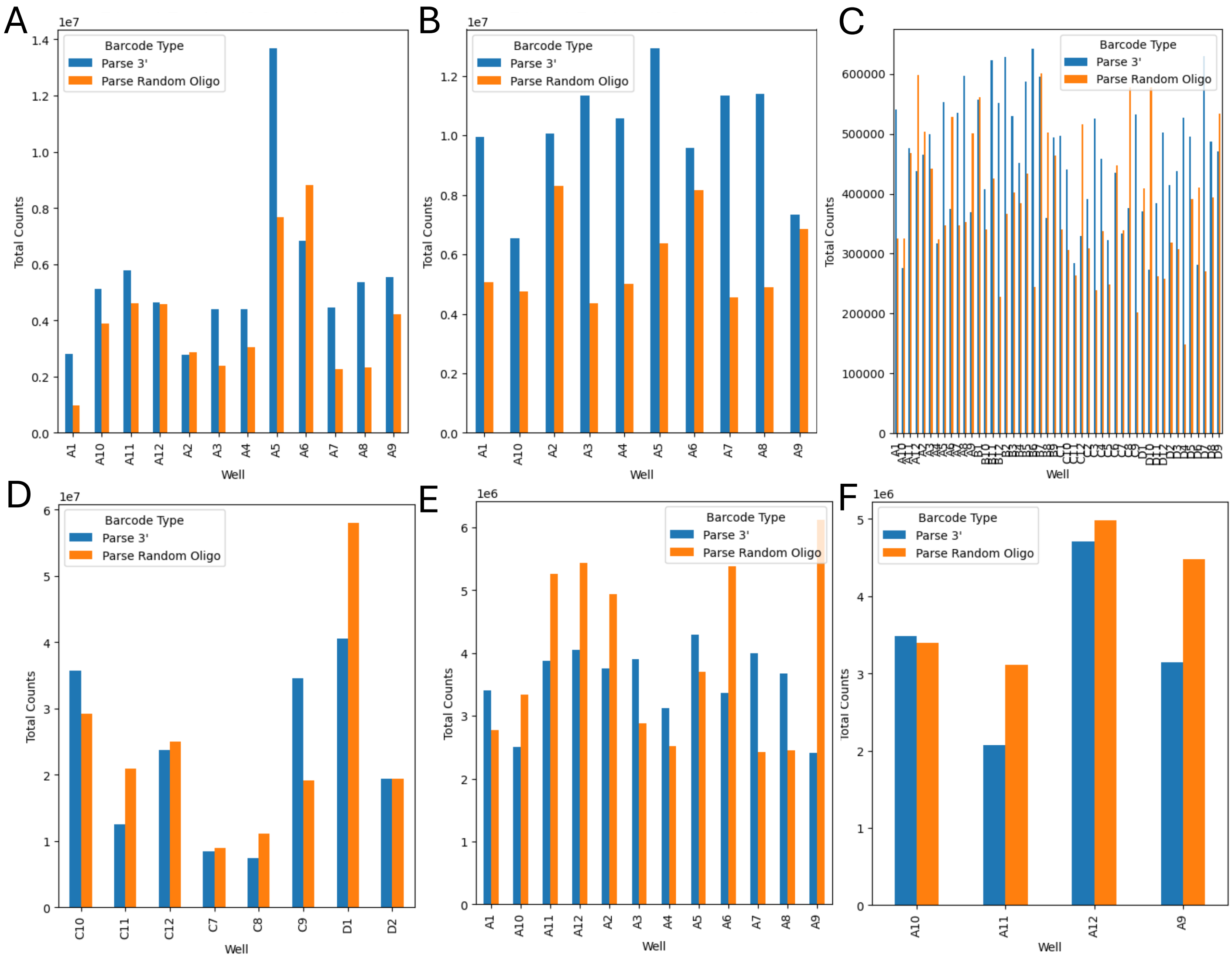


**Fig S12.** (A) Bar plot of the number of oligo(dT)- and random-hexamer-primed counts originating from each Parse Evercode WT round 1 well, for Exp 1. (B–G) Same as (A), but for Exp 2, Exp 3, Exp 4, Exp 5a, Exp 5b, and Exp 6 respectively.
